# VN1K: a genome graph-based and function-driven multi-omics and phenomics resource for the Vietnamese population

**DOI:** 10.1101/2025.04.15.648991

**Authors:** Trang T. H. Tran, Tham H. Hoang, Mai H. Tran, Nguyen T. Nguyen, Dat T. Nguyen, Tien M. Pham, Nam N. Nguyen, Giang M. Vu, Vinh C. Duong, Quang T. Vu, Thien K. Nguyen, Sang V. Nguyen, Hien Q. Vu, Trang M. Nguyen, Toan Dang, Hoang Nguyen, Tuan Do, Cuong Le, Hung T. T. Nguyen, Nam Q. Le, Quang-Huy Nguyen, Linh T. Le, Thang Pham, Minh Dao, Duc M. Vu, Huong T. T. Le, Tho D. Ngo, Liem T. Nguyen, Yen Hoang, Dat X. Dao, Giang H. Phan, Loan Nguyen, Chi Trung Ha, Hung N. Luu, Vinh Le, Thinh Tran, Ly Le, Nguyen Thuy Duong, Duc-Hau Le, Quan Nguyen, Van H. Vu, Nam S. Vo

**Affiliations:** Center for Biomedical Informatics, Vingroup Big Data Institute, Hanoi, Vietnam; R&D Department, GeneStory JSC, Hanoi, Vietnam; Asian School of the Environment, Nanyang Technological University, Singapore; Hi-tech Center, Vinmec Healthcare System, Hanoi, Vietnam; Vinmec Research Institute of Stem Cells and Gene Technology, Vinmec Healthcare System, Hanoi, Vietnam; Nursing Division, Vinmec Healthcare System, Hanoi, Vietnam; Hanoi Medical University, Hanoi, Vietnam; Institute for Molecular Bioscience, The University of Queensland, Brisbane, QLD 4072, Australia; UPMC Hillman Cancer Center, University of Pittsburgh, Pittsburgh, PA 15232, USA; Department of Epidemiology, School of Public Health, University of Pittsburgh, Pittsburgh, PA 15261, USA; University of Engineering and Technology, Vietnam National University Hanoi, Hanoi, Vietnam; Institute of Genome Research, Vietnam Academy of Science and Technology, Hanoi, Vietnam; School of Information and Communications Technology, Hanoi University of Science and Technology, Hanoi, Vietnam

## Abstract

Vietnam, the 16th most populated nation, remains profoundly underrepresented in global genomic databases. Here, we present VN1K, a first-ever comprehensive and well-curated resource of multi-omics data with a wide-range of phenotypic information of 1,011 unrelated Vietnamese individuals. High-depth short-read whole-genome sequencing data were generated for all samples along with various - omic data, including microarray, long-read whole-genome sequencing, and RNA sequencing. Using a high-sensitivity variant detection pipeline, which included a pangenome graph reference and a deep-learning framework, we identified nearly 40 million variants of which 8.5 million are novel with nearly 900 thousand short insertions/deletions and 39 thousand structural variants. Specifically, VN1K featured a first-ever whole-genome methylation profile based on long read sequencing. A genotype imputation panel was also created with the highest accuracy on the Vietnamese population. Variants with significantly different allele frequencies in the Vietnamese population compared to others were found to be functionally significant, especially in genes associated with immune diseases (*HLA-B, KIR3DL3, KIR2DL1, KIR2DL4*) or drug responses (*CYP2C19, CYP2D6, VKORC1, CYP2B6*). We were also able to map various loci related to hepatitis B virus infection as well as six disease traits, including triglyceride levels, LDL-C, serum glucose levels, HbA1c, and levels of two liver enzymes (ALT and AST). VN1K dataset is accessible via genome.vinbigdata.org, an integrated platform with both linear and graph-based genome browser for facilitating data exploration, research, and applications in precision medicine.

## INTRODUCTION

Vietnam is profoundly underrepresented in the current global genomic databases despite the large population and extensive genetic and phenotypic diversity. As of December 2024, Vietnam has a population of approximately 100 million people and is ranked first in the mainland region of Southeast Asia and the 16^th^ worldwide with a long and complex immigratory history, resulting in extensive ethnolinguistic and genetic diversity (Liu et al. 2020). As Vietnam is located in a key geopolitical region connecting South, East, and Southeast Asia, understanding its genetic structure is critical to further elucidate the genetic diversity of the larger neighboring areas. Despite these facts, the current widely-used whole genome sequencing (WGS) dataset for Vietnamese comprises only 99 Kinh in Ho Chi Minh City, Vietnam (KHV) samples in the 1000 Genomes Project (1kGP-KHV) (1000 Genomes Project Consortium et al. 2015), severely under-representing the approximately 100 million people across the 64 provinces. Recent WGS projects for the Vietnamese population were performed with just either a few samples with high coverage (a trio at 30X) (Hai et al. 2015) or few hundreds of samples with low-to-medium coverage (99 WGS at 4x from 1kGP-KHV, 105 WGS at 15x and 200 WES from in-house data) (Le et al. 2019). In recent Asian genome projects, including the GenomeAsia100K Pilot Project (GAsP) (GenomeAsia100K Consortium 2019), in which Southeast Asian samples account for nearly 20%, the genetic data from the Vietnamese population is not present. This lack of knowledge highlights the underrepresentation of Vietnamese population in the global genomics, possibly due to a long-term exclusion from Vietnamese origins from the database.

Despite the remarkable expansion of genome projects, current studies have mostly focused on European individuals (or Caucasian ancestry), which capture around 80% of participants in genome-wide association studies (GWAS) (Popejoy and Fullerton 2016). Asians, accounting for 60% worldwide population with diverse ethnicities (20007900), are still underrepresented in large databases such as gnomAD (Karczewski et al. 2020) or TOPMed (Taliun et al. 2021). Although some groups such as South Asians and East Asians have been thoroughly studied in the 1kGP, many of them are often lumped together in large projects. Currently, an increasing number of Asian genome projects have been conducted, leading to the robust discovery of novel variants compared to European-centric genomic datasets (Nagai et al. 2017) (Jeon et al. 2020) (Wu et al. 2019) (Kutanan et al. 2021). Notably, Southeast Asian genomes such as Malaysian in SG10K yields more novel variants than both South and East Asian (Wu et al. 2019). This indicates that Southeast Asia, which accounts for 8.5% of the world’s population and has a long and complex history of human migration within the region and from South and East Asia regions, is greatly understudied and requires more GWAS studies. Inclusion of the Southeast Asian genomes in the genomic databases also significantly reduced the error rates of genomic imputation for ethnically related populations such as Melanesians and Papuans from Oceania. Further investigation of the genetics of Southeast Asians, including Vietnamese, will undoubtedly uncover rare and population-specific variants, inform variant prevalence, and provide a comprehensive reference genome.

Here we present VN1K, the first large-scale genome program for the Vietnamese population (1,011 individuals) with high-depth WGS (37X median, 30X for 98% samples) coupled with latest genomic sequencing technologies and functional assays (PacBio and Oxford Nanopore Technologies long-read sequencing, Axiom™ Precision Medicine Diversity Array (APMDA) genotyping, and Illumina RNA sequencing) along with a wide-range of common phenotypic traits (*n*=34 in total, including demographic, anthropometric, biochemical, and clinical traits). This pioneering study presents the most comprehensive dataset with a significant number of novel variants detected, stringently curated and functionally annotated to investigate their phenotypic and potential clinical impacts.

First, we identified a large number of nearly 8.5 million novel variants, allowing us to not only unravel the genetic diversity of the Vietnam population, but also create an imputation reference for Vietnamese people with high accuracy. Second, we constructed a pangenome graph based on genomic data of multiple long-read and short-read platforms, which could enhance the ability to detect structural variants. Third, all genomic variants (SNPs, INDELs, and SVs) were annotated, followed by focused investigation of the pathogenic variants with considerable frequencies in the population, aiming to improve genetic screening panels for various clinical applications. Fourth, we analyzed the genotypes of several important pharmacogenes and immunogenes to investigate the unique genetic patterns of Vietnamese genomes. Fifth, and for the first time, this study reported methylation profiles, adding an important layer of the functional genome, beyond DNA sequence, that is the epigenetics regulation at the genome-wide scale. This was stringently acquired from combining results of two long-read sequencing technologies, serving as the foundation to refine the putative effects of DNA variants within or outside the active regions of the genome. Sixth, we carried out Genome Wide Association (GWA) analyses for 12 collected clinical phenotypes and showed that the VN1K resource, although still limited in sample size, is useful to map genotype-clinical phenotype associations in the Vietnamese population. Seventh, we refined the genetic structure and evolution of the Vietnamese population, identifying differences between Vietnamese regions and other populations. Finally, to put it all together, a big data platform called MASH (Management, Analysis, Sharing, and Harmonization) was implemented to enable management, analysis, and sharing of VN1K dataset. MASH will also facilitate global collaboration to explore and integrate with the Vietnamese genomic data. Overall, this study provided the much needed VN1K data resource with the most comprehensive information about the genotypes and phenotypes of the Vietnamese population, providing a high-quality resource for genetic and biomedical research on Vietnamese and related populations of Asian-origin, and contributes significantly to research and applications in the field across Southeast Asia, Asia, and the world.

## RESULTS

### 1. VN1K reveals a largest-ever number of novel variants and improves imputation accuracy

VN1K comprises the largest collection of Vietnamese WGS samples, with 1,011 individuals geographically representative for 54 of the total 63 provinces and cities of Vietnam (**Figure 1a**). Our stringent pipeline defined the final VN1K genomic variant callset with 39,712,388 variants, consisting of 34,875,006 bi-allelic SNPs, 193,553 multi-allelic SNPs, 5,111,833 bi-allelic INDELs, and 1,433,546 multiallelic complex sites. Furthermore, the largest number of CNVs was identified with a total of 21,255, consisting of 17,192 deletions (DEL) and 4,063 duplications (DUP) (**Suppl. Figure 1** and **Suppl. Table 1**). Compared to SNPs from 1000 Genomes Project high coverage (1kGP-HC), Genome Aggregation Database (gnomAD v3) and Database of Single Nucleotide Polymorphisms (dbSNP151), we mapped as many as 8,480,031 novel variants, likely specific for the Vietnamese and other under-represented populations of Asian-origin. Among those novel variants, 7,584,284 are SNPs and 895,747 are INDELs, most of which were rare (allele count > 2 and allele frequency (AF) ≤ 0.01) and intergenic/intronic (**Figure 1b**). Many of these specific variants are functionally relevant. For instance, we found 166 human leukocyte antigen (HLA) class I and II alleles, of which 79 were novel (**Figure 1c**). To further investigate the VN1K novel variants, we compared them to the most recent Vietnamese genomic database which included a collection of 406 individuals, comprising105 whole genomes, 200 whole exomes, and 101 published whole genome samples. Strikingly, our results showed that the vast majority (95.75%) of VN1K novel variants are not in this database. This unique and important addition was not only derived from a larger number of 1011 samples, but more importantly, resulted from well-curated, high-depth sequencing (30-60X), and multi-platform sequencing data, including short-read data from Illumina, and long-read data from PacBio and Oxford Nanopore Technologies (ONT) platforms. The large number of new variants detected by the VN1K enabled more accurate imputation for the Vietnamese population than any other reference panels (**Figures 1d-e**). In particular, the merged panel VN1K_SG10K has the best performance in imputing the genotype of the Vietnamese samples, which was validated by our Affymetrix PMRA microarray data of 96 randomly selected samples. VN1K also led to the highest aggregate R^2^ (i.e., the coefficient of determination) between the imputed and true genotypes across all bins of minor allele frequency (MAF) (**Figure 1d**). Our imputation panel has remarkable translation implications, for example, in HLA imputation where the combined panel VN1K_1kGP-HC and the VN1K provided comparable accuracy for both HLA class I and II genes which outperformed 1kGP-HC alone (**Figure 1e**). Therefore, we expect that the custom imputation panel could greatly benefit GWAS on the Vietnamese population by using SNP arrays followe by Vietnamese-specific imputation. Indeed, we have recently assessed the application of imputation on estimating polygenic risk scores (Pham et al. 2022).

**Figure 1.**
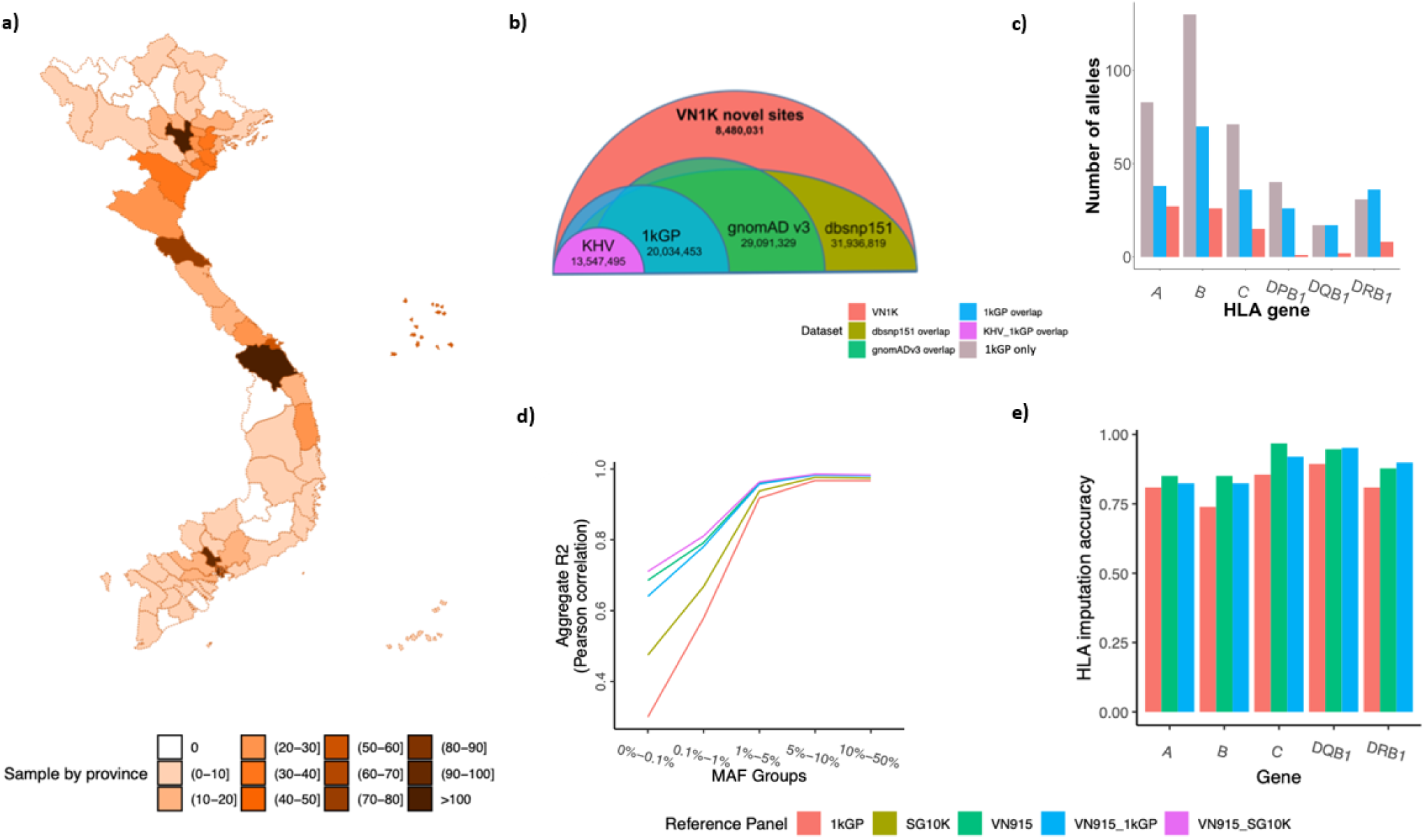
Sample collection, statistical analysis of novel sites, and the application of SNP-HLA imputation techniques to the VN1K dataset. The 1,011 unrelated Vietnamese individuals across 54/63 provinces and cities revealed millions of novel variants not reported in existing databases. **a**) Geography of collected samples. 1,011 Vietnamese individuals declared as Kinh-ethnic for at least 3 generations and confirmed as healthy by their general health check information. Participants came from 54 of the total 63 administrative divisions of Vietnam. **b**) The total number of variants in VN1K overlaps with 1kGP-HC, gnomAD v3 and dbsnp151. **c**) The number of HLA alleles class I and II in 1kGP-HC and novel alleles in VN1K. d) Imputation accuracy measured by aggregated R2 of all imputed variants in different MAF bins for Vietnamese-Kinh population based on 3 single panels (1kGP-HC, SG10K, VN1K) and 2 merge panels (VN1K+1kGP-HC and VN1K+SG10K). **e**) HLA imputation accuracy comparison between different imputation panels. Abbreviation: gnomAD: The Genome Aggregation Database; dbsnp151: Single Nucleotide Polymorphism Database version 151; 1kGP: 1000 Genomes Project; 1kGP-HC: 1000 Genomes Project – High Coverage; SG10K: Singaporean Genomes Project.

### 2. Pangenome graph reference combined with a deep-learning framework improves structural variant detection

As this project aimed to produce the most extensive and stringent collection of SVs, we combined and cross-validated SVs from the short read Illumina data with SVs from multi-platform long-read sequencing data (PacBio and ONT). To best represent and enable broad usage of such a comprehensive variant resource for SNPs and SVs, we built a pangenome graph reference, our main reference and hereby referred to as the Vietnamese Genome Reference (VGR). VGR integrated into graph the assemblies of two VN1K samples, deeply sequenced by long-read PacBio and ONT platforms (one male and one female, **Suppl. Table 2**). These assemblies covered up to a total length of 2.76 Gbp with N50 and L50 of up to 31,298,235 and 25, respectively. To improve the comprehensiveness and cross-database compatibility with public resources, we also incorporated assemblies of three KHV samples from the Human Pangenome Reference Consortium (HPRC) database into the VGR as an additional track. Combined with a deep-learning framework, which utilized the Convolutional Neural Network model (CNN) to refine the initial result from existing tools for SV calling, VGR enabled accurate detection of population-specific SVs, which were difficult to obtain using just the linear reference GRCh38. First, using GRCh38, we detected 2,848±362, 5,046±285, 666±74, 61±11 (mean±sd) INS, DEL, DUP, and ITX, respectively (**Figures 2a-b**), of which the overlapping (short-read vs. long-read) between DEL was higher than that of INS (**Figure 2c**). Interestingly, we found that Vietnamese and East Asian (EAS) superpopulation shared much more common SVs than with the 1kGP, with the overlapping rates of 76.2% DEL and 83.9% INS (**Figure 2d**). By examining a set of SVs that were not presented in 1kGP-HC (represented by the blue area in **Figure 2d**), we found a significant SV subset with high AF, which potentially represented the SVs specific to the Vietnamese population. Furthermore, we found that VGR significantly improved the SV calling compared to using GRCh38, for both short-read and long-read data. As exemplified in one randomly selected sample, 9,343 and 7,816 were detected with VGR compared to 8,774 and 3,762 with GRCh38 using long-read data and short-read data (**Suppl. Figure 2**). The ability to capture the diverse and population-specific SVs through a graph-based pangenome reference and deep learning framework could unlock new insights into the genomic landscape, detailed in **Methods** session and **Suppl. Methods.**

**Figure 2.**
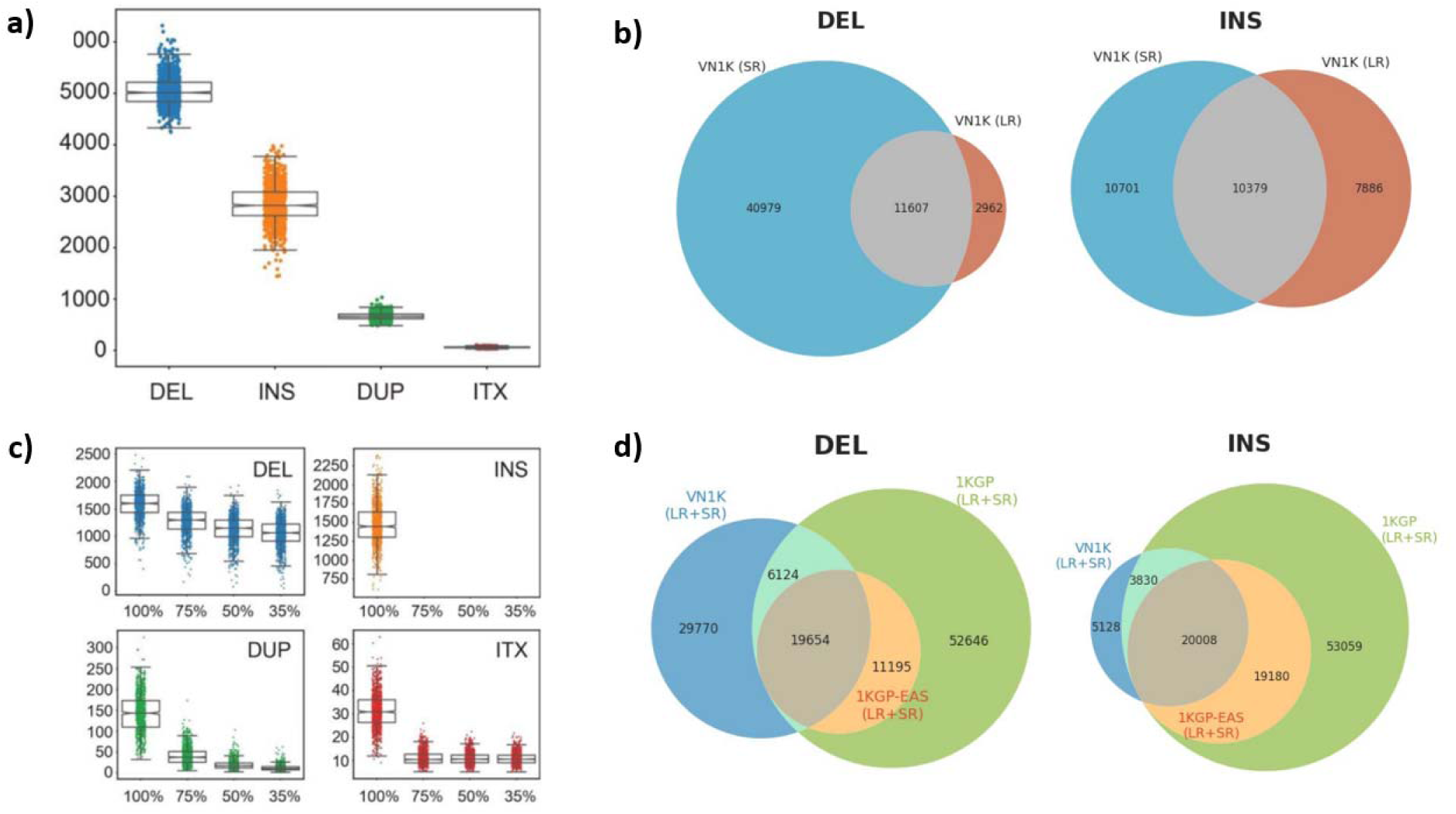
SV profiles for VN1K with short-read (by Illumina) and long-read (by PacBio) sequencing. The SV profiles indicated that utilizing long-read sequencing approaches significantly improved coverage for the Vietnamese population, and other populations also exhibit similar SV profiles of 1kGP-HC. a) The distribution of detected SVs of DEL, INS, DUP, ITX; b) SV sites compared within VN1K for short-read and long-read results; c) each SV type requiring reciprocal overlaps at varying levels 100%, 75%, 50%, and 35% between SVs, only 100% overlap was shown for INS. d) SV sites compared between VN1K and 1kGP-HC databases. Abbreviation: SR: short reads, LR: long reads, EAS: East Asian, SV: structural variant, ITX: Intra-chromosomal translocation, DUP: Duplication, NS: Insertion, DEL: Deletion.

### 3. Functional genomic annotation analysis and implications in genetic disease screening

To investigate rare mutations, doubletons, and singletons that are unique to the Vietnamese population, we categorized the SNPs, INDELs, and SVs in VN1K by AF and compared them to cu rent popular databases, including 1kGP-HC, gnomAD v3, and Centers for Common Disease Genomics (CCDG) (**Figure 3a**). Of all novel variants, intronic and intergenic variant regions accounted for 54.3% and 29.6%, respectively. There were only 0.95% of variants in the coding and splicing regions (**Figure 3b**). We defined an important SV as the one that overlaps with multiple regions of a gene, the region at the higher level of the functional hierarchy (in the order of CDS/exon, UTR, intron, flanking). Two ranges of flanking length were specified by default (i.e., 5Kb and 50Kb). The SVs in intronic and exonic regions attributed to 59.2% and 5.29%, respectively (**Figure 3c**). Importantly, we found that nearly half of the novel single nucleotide variants (SNV) were missense, 22.06% were splicing, and 21.72% were synonymous (**Suppl. Figure 3**). The VN1K database enabled the discovery of specific variants in disease-associated genes such as *GCNT2* and *PMFBP1*, which exhibited a high prevalence (14.84%) in the Vietnamese population but rare (less than 0.5%) in other large populations of European. In contrast, the pathogenic variants associated with *HNRNPUL2* and *BSCL2*, which linked to congenital generalized lipodystrophy and breast carcinoma, were less prevalent in either the Vietnamese and East Asian (EAS) populations (less than 10% and 11.9%, respectively), yet more common in other populations (e.g., 28.93% in Europe and 28.12% in South Asia in 1kGP-HC). Notably, *PMFBP1*, *MTSS2*, *C9, SRD5A2*, and *HFE,* which were not featured in the American College of Medical Genetics and Genomics (or ACMG) prenatal screening for genetic disorders, harbored known pathogenic variants with considerable AF (ranging from 1.48% to 3.26%) in the Vietnamese population (**Suppl. Figure 4**).

**Figure 3.**
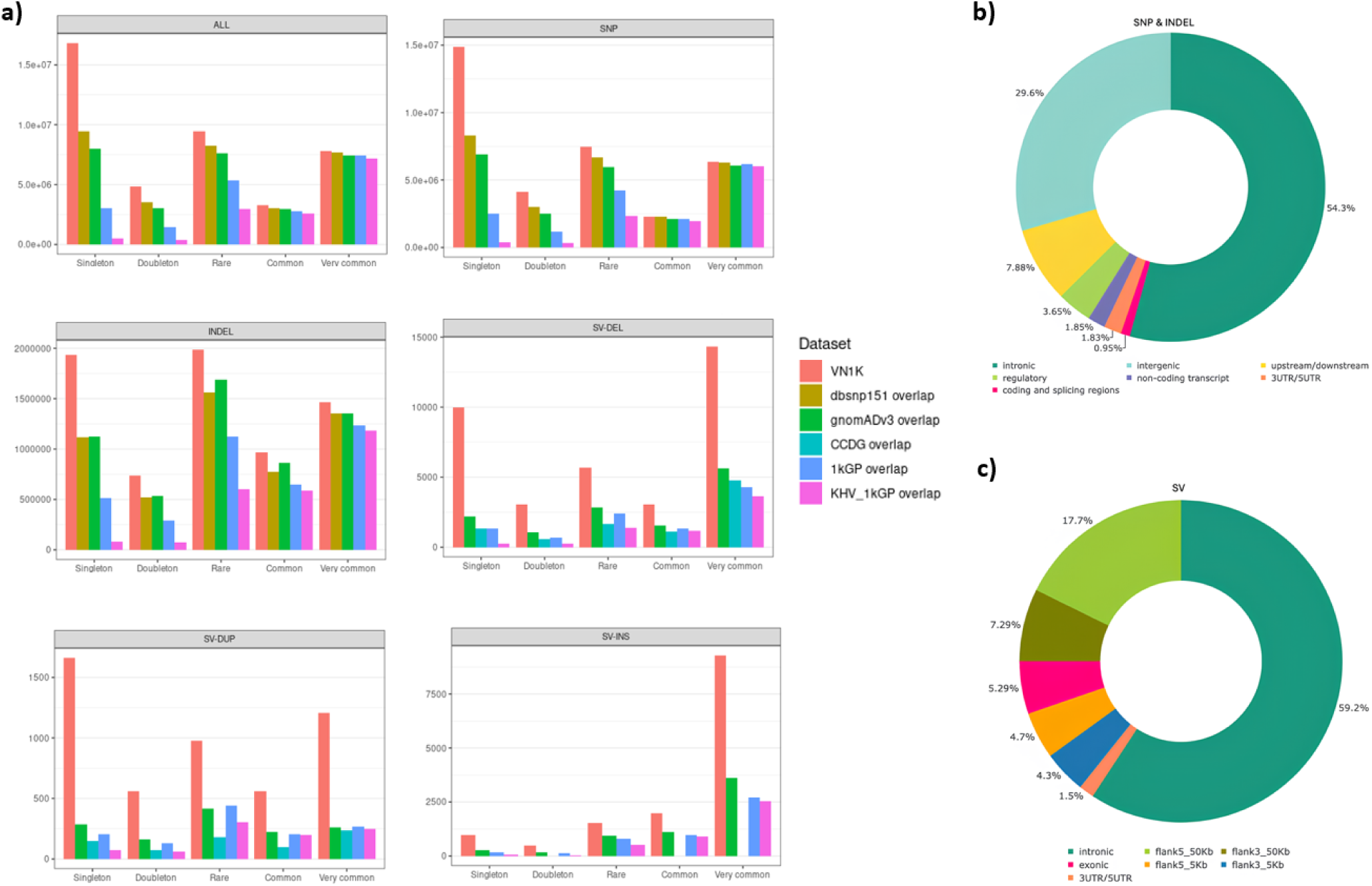
Functional genome annotation of the unique variants for the Vietnamese population. (a) Number of SNPs, INDELs, and SVs of VN1K compared to the public databases. We found that the mutational profiles relative to the public databases unveil rare mutations, doubletons, and singletons unique to the VN1K dataset. All variants were categorized by allele frequency and by comparison to the 1kGP-HC, gnomAD v3, and dbsnp151 dataset. The SVs were categorized by allele frequency and by comparison to the 1kGP-HC, gnomAD v4 and CCDG. (b-c) Summary of the variant consequences based on annotation of the VN1K’s novel callset (SNPs & INDELs, SVs). Abbreviation: CCDG: Centers for Common Disease Genomics.

### 4. The immunogenomics and pharmacogenomics references for the Vietnamese population

The VN1K genomic variant data also allows us to assess the genetics of the Vietnamese immune system and their potential implications in drug responses, a pioneering contribution towards precision medicine in an under-represented country. The interplay between HLA and killer cell immunoglobulin-like receptors (KIR) were known to have significant impacts on immune function and disease susceptibility (Y. Li et al. 2021) (Tham H. Hoang et al. 2022) (Peng et al. 2021). First, to assess the generalizability of our data, we compared allele frequency in different ethnicities, and found that the HLA’s AF exhibited consistent patterns with most public datasets of the Kinh ethnicity (e.g., KHV individuals in 1kGP-HC) but varied significantly across different ethnicities in Vietnam (e.g., Muong in Hoabinh province from Allele Frequency Net Database (Gonzalez-Galarza et al. 2018)) (**Figure 4a**). *HLA-B*15:02*, which is highly associated with many Adverse Drug Reactions (ADR) such as carbamazepine-induced SJS/TEN, was the most common allele, accounting for 19.05% of HLA-B allelic diversity. *HLA-B*15:02* also represented a significantly higher AF (23.22%) than those observed in other populations of 1kGP-HC (e.g., 6.17% in EAS, 8.38% in SAS of 1kGP-HC). Allele and gene frequencies of KIR genes showed that all four framework genes (i.e., *KIR3DL3*, *KIR2DL1*, *KIR2DL4*, and *KIR2DL2*) and one telomeric gene (i.e., *KIR3DL1*) were most frequently present in on gene-content haplotypes of the Vietnamese population (**Figure 4b**).

**Figure 4.**
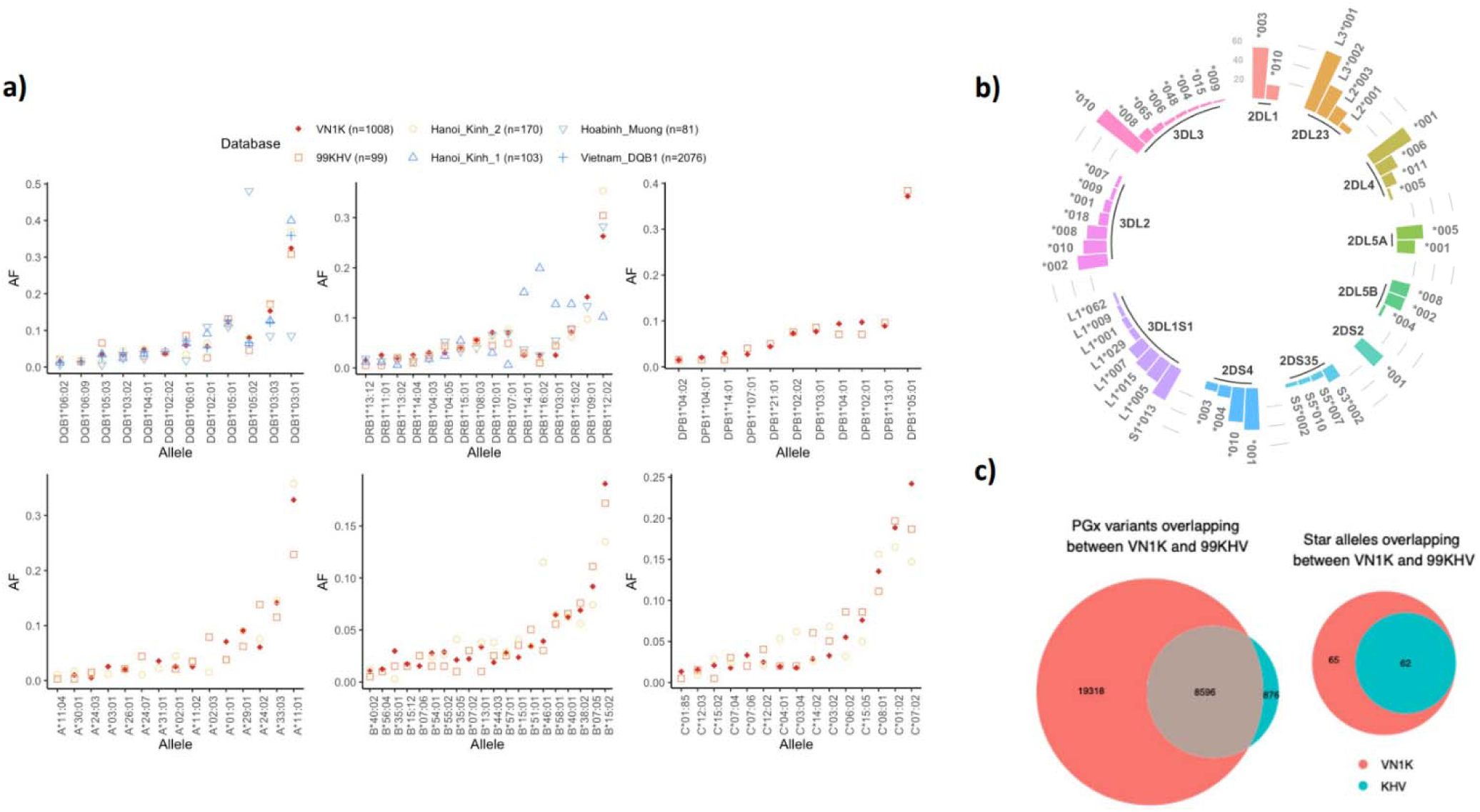
The immunogenomics and pharmacogenomics profiling of VN1K. A comparative study involving the VN1K and other existing datasets from Vietnam examined actionable genotypes and diplotypes. a) The AFs of six *HLA* genes in the Vietnamese population from VN1K, KHV and public data from allelefrequencies.net. Note that the Vietnam_DQB1 (n=2076) obtained the result from genotyping data of gene panel testing. b) The AFs of KIR genes called from the VN1K dataset. c) The number of PGx variants and star allele calls from VN1K and KHV datasets. Abbreviation: KHV: Kinh in Ho Chi Minh City, Vietnam (The 1000 Genomes Project).

Another clinically important and complex genomic region is pharmacogenes (PGx), which affect the drug properties (i.e., pharmacokinetics and pharmacodynamics) (Rollinson, Turner, and Pirmohamed 2020). VN1K included a total of 27,914 variants on the PGx genes, which were further assigned to 127-star alleles (or gene-level haplotype patterns that related to drug metabolisms) (**Figure 4c**). For instance, the *CYP2D6*10* allele has an AF of 23.16%, suggesting a key genetic contribution to the observation that more than half of the Vietnamese carry the phenotype of intermediate metabolizer (IM). Two non-functional *CYP2C19* alleles, *\*2* and *\*3*, have AF of 26.37% and 5.21%, respectively. In contrast, *CYP2C19*17,* a more functional allele that ís associated with ultra-rapid metabolism of drugs, has an AF of only 1.09%. Furthermore, we found that the 99 KHV samples in the 1kGP-HC dataset covered only 9,472 variants assigned to 65 star-alleles, all of which were detected in our VN1K reference, suggesting the comprehensiveness of our dataset in PGx gene regions. Specifically, top genes with highest Jost’s D genetic distance between VN1K and 1kGP-HC (excluding KHV) were *VKORC1*, *CYP2D6*, *CYP2B6*, followed by *NAT2*, and *SLCO1B1* (**Suppl. Figure 5**). Taken together, our analysis presents the most comprehensive characterization of these important genomics regions, highlighting variant AF unique to Vietnamese and other ethnic groups.

### 5. Whole-genome methylation analyses reveal specific and functionally relevant epigenomic signatures

We posited that functional features of a reference genome are not only dependent on DNA sequences but also on the epigenomic states. For the first time, we built a Vietnamese epigenomic reference starting with deep methylation profiling of two individuals (i.e., one male and one female) and 4 pools of 40 individuals using both PacBio and ONT technologies, followed by an integrative analysis of RNA-seq data (Mayne, Berry, and Jarman 2021). We identified 27,605,743 highly confident methylation sites, representing approximately 95% and 98% of the sites from PacBio and ONT, respectively. The two results were consistent across all chromosomes, with Pearson correlation coefficient (R) of 0.922 **(Figure 5a)**, although we noted that ONT data appeared to show more high-confidence sites (both methylated and unmethylated), whereas PacBio data showed more low-methylated and inter-methylated sites **(Figure 5c)**. Nevertheless, the consensus sites enabled us to investigate not only methylation patterns within pharmacogenomic regions for Vietnamese but also differentially methylated regions (DMRs) between Vietnamese and other ancestries of EUR and AFR, corroborating an additional layer of genomic diversity **(Figure 5b)**. We found DMRs within clinically relevant genes that are higher or lower in the Vietnamese samples compared to the European samples, as illustrated by the locus plot generated from Methylartist, where the similar score trends between the two sexes are found higher in the Vietnamese population **(Figure 5d)**. The comparison of expression of genes located in DMRs with the *GAPDH* gene as a control showed that the top differentially methylated genes, *DLGAP2,* also exhibited a lower gene expression level in the Vietnamese dataset compared to the European data, consistent with the results showing hypermethylation in Vietnamese and hypomethylation in European (*P*<0.15 vs. *P*<9.22e-5) **(Figure 5e)**. In such locations, the SNP effect sizes generally estimated at the population scale in Caucasian would be different to that in the Vietnamese due to different states of the genome, which may not be transcribed or accessible for transcription factor binding, even if a SNP presents. An analysis of 14 samples using RNA-Seq data for the genes associated with DMRs further confirmed the unique epigenetic signatures and consequences on gene expression. Many of those genes are enriched for important phenotypes such as alcohol drinks (*P*<6e-5), triglyceride metabolisms (*P*<1e-3), fatty lipid metabolism (*P*<2e-4), and virus perturbation (*P*<2e-2) **(Figure 5f)**. Taken together, such epigenetic and transcriptomic associated pathways suggest potential mechanisms for the observed higher incidence in liver-related diseases (Duong et al. 2009) (Tung et al. 2022)

**Figure 5.**
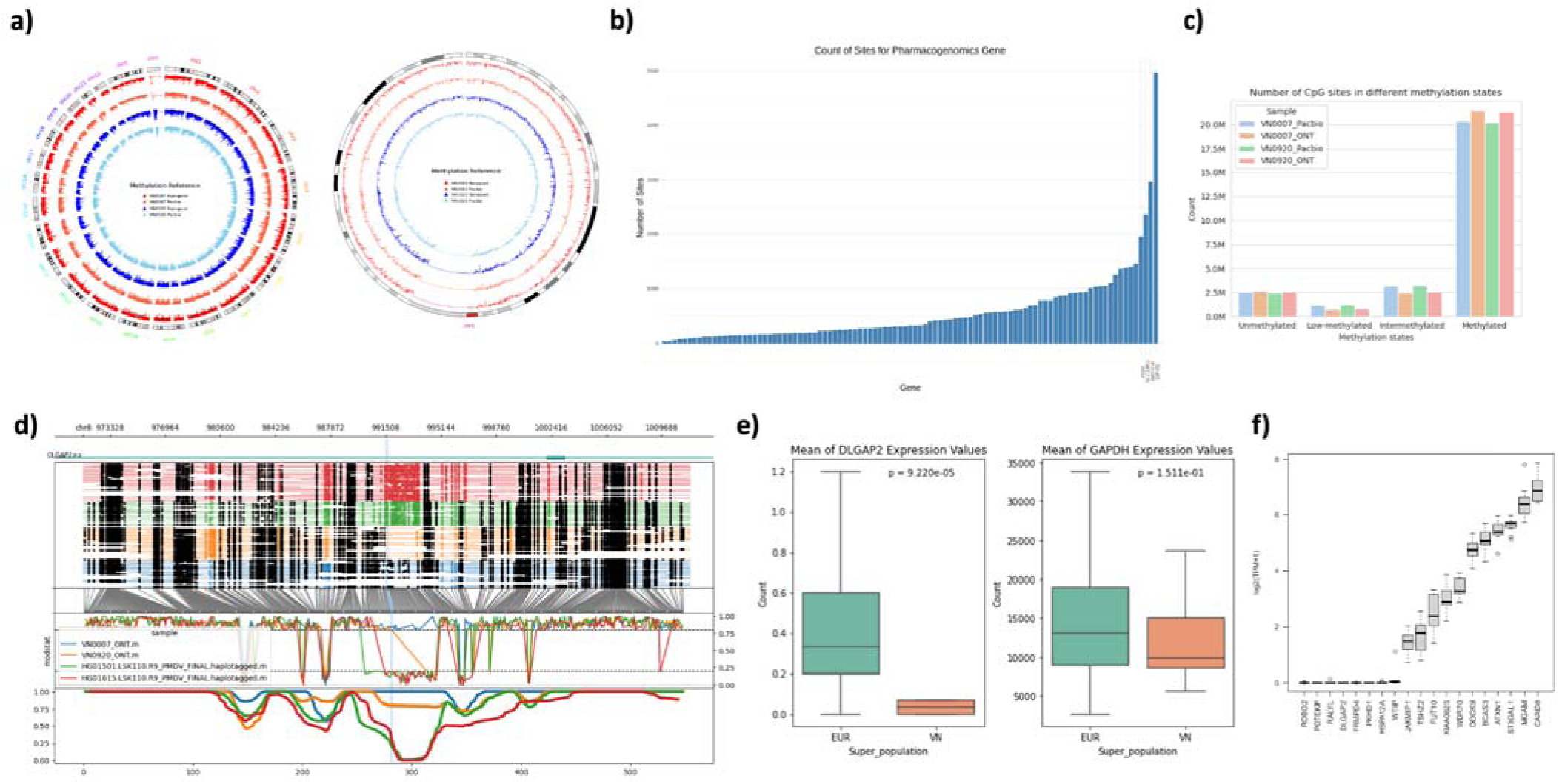
The consensus methylation reference for Vietnamese individuals. (a) The circos plot of the Vietnamese methylation reference panel for all chromosomes (1-22, X, Y) and the zoom in chromosome 1. (b) Consensus of CpG sites of Pacbio and ONT on 100 pharmacogenomic genes in PKSeq panel developed by RIKEN. (c) The distribution of methylation levels between Pacbio and ONT was categorized into four distinct groups. (d) The locus plot generated from Methylartist shows the differentially methylated regions (DMRs) between Vietnamese and European individuals were compared in gene *DLGAP2*. (e) The box plot compares the mean gene expression of genes located in differentially methylated regions (DMRs) to the expression of the *GAPDH* gene. (f) The methylation and RNA-seq gene expression of log2(TPM+1) was shown in the bottom panel of the figure. The selected gene set is enriched in alcohol drinking, lipids and other liver induced dataset.

### 6. Multi-phenotypic traits and metagenomics analyses identify functional loci of interest

We further investigated the interplay between phenotypic traits and metagenomics of VN1K. First, an association analysis was performed with 12 quantitative clinical traits and 8,536,859 variants from 810 samples with full trait information using additive genetic models (**Suppl. Table 3**). Despit the relatively small sample size, we found several independent loci surpassing the GWAS threshold (*P*<5.86e-9, Bonferroni-corrected) (**Figure 6**). These loci were significantly associated with six traits, including 1) triglyceride levels, 2) low-density lipoprotein cholesterol (LDL-C), 3) serum glucose levels, 4) *HbA1c*, and 5 and 6) the levels of two liver enzymes (i.e., alanine aminotransferase-ALT and aspartate aminotransferase-AST). These results were then further refined by mapping GWAS catalog data to VN1K and comparing the VN1K common variants with those from other ethnic groups.

**Figure 6.**
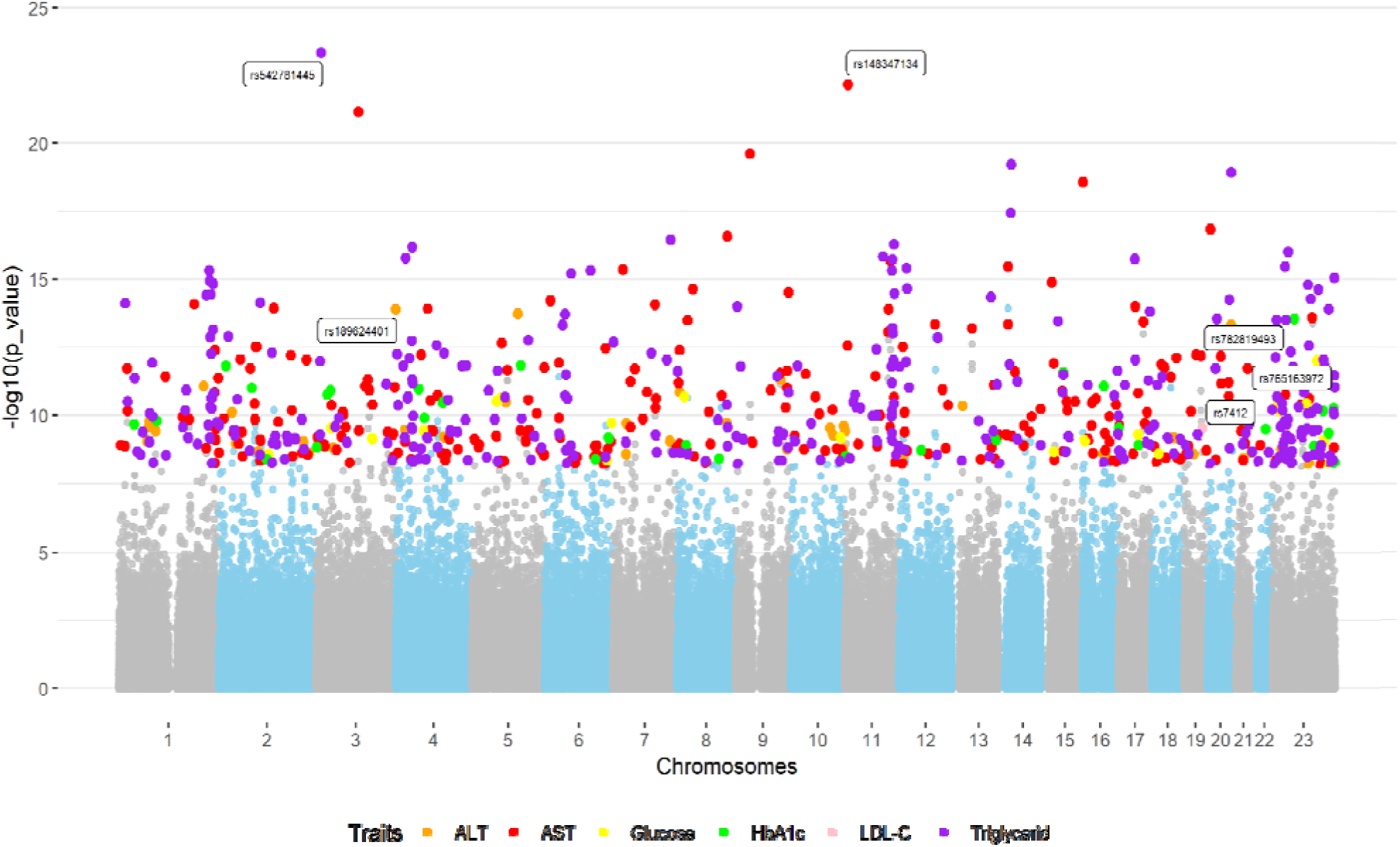
Multi-trait association analysis. 12 quantitative clinical traits and 8,536,859 variants from 810 samples in the VN1K dataset using additive genetic models. Manhattan plot revealed several independent loci surpassing the GWAS threshold (*P*<5.86*e-9, Bonferroni-corrected significance). These loci were associated with six traits: triglyceride levels, LDL-C, serum glucose levels, Hemoglobin A1c, and levels of two liver enzymes (ALT/AST).

Second, we examined the presence and possible effects of common viruses in the ge etic structure of the Vietnamese population. We found eight viruses with genetic material mapped to VN1K genomes, in which hepatitis B virus (HBV) presented with the highest frequency (99 samples, or 9.8%), having mean read depth over 10 (**Suppl. Figure 6**). These results, for the first time, provided a genetic-based assessment for the HBV epidemic in Vietnam, which is among the top 11 countries globally suffering from chronic viral hepatitis, with the prevalence of hepatitis B surface antigen (HBsAg) over 8% (Zampino et al. 2015). Such findings consistent between genetics data and epidemiology data warranted future validation studies, coupled with results from the epigenetic and transcriptomic analysis.

### 7. VN1K suggests refining genetic structure and evolution of the Vietnamese population

Vietnam was known to have a long and complex immigratory history with extensive ethnolinguistic and genetic diversity (Liu et al. 2020). This was examined more thoroughly in our ADMIXTURE analysis on the VN1K data combined with WGS data from the Human Genome Diversity Project (HGDP) (**Figure 7**). With K=10 for ADMIXTURE analysis, our results showed that Vietnamese shared components with the Chinese Dai, Singapore-Malaysian, and Japanese. Notably, Group 1 and 2 (“group’ referring to ‘genetic cluster’), major components for the Dai population (Chinese Dai in Xishuangbanna, China [CDX]) and the VN1K population, are specific to the East Asia Region. VN1K also aligned closely with CDX, consistent with previous reports (Tham Hong Hoang et al. 2024) (Nguyen et al. 2024). The genetic diversity was further highlighted by FSt values between Vietnam and other Asian groups in the HGDP. Our analysis showed a trend for the higher degrees of genetic difference of Kinh people for Central Asian groups than for East Asian groups (**Suppl. Figure 7**). The notable difference in genetic distance as shown in our analysis suggests the need to include Vietnamese data in multi-ancestry analyses, especially among those ethnicities that have not been well studied.

**Figure 7.**
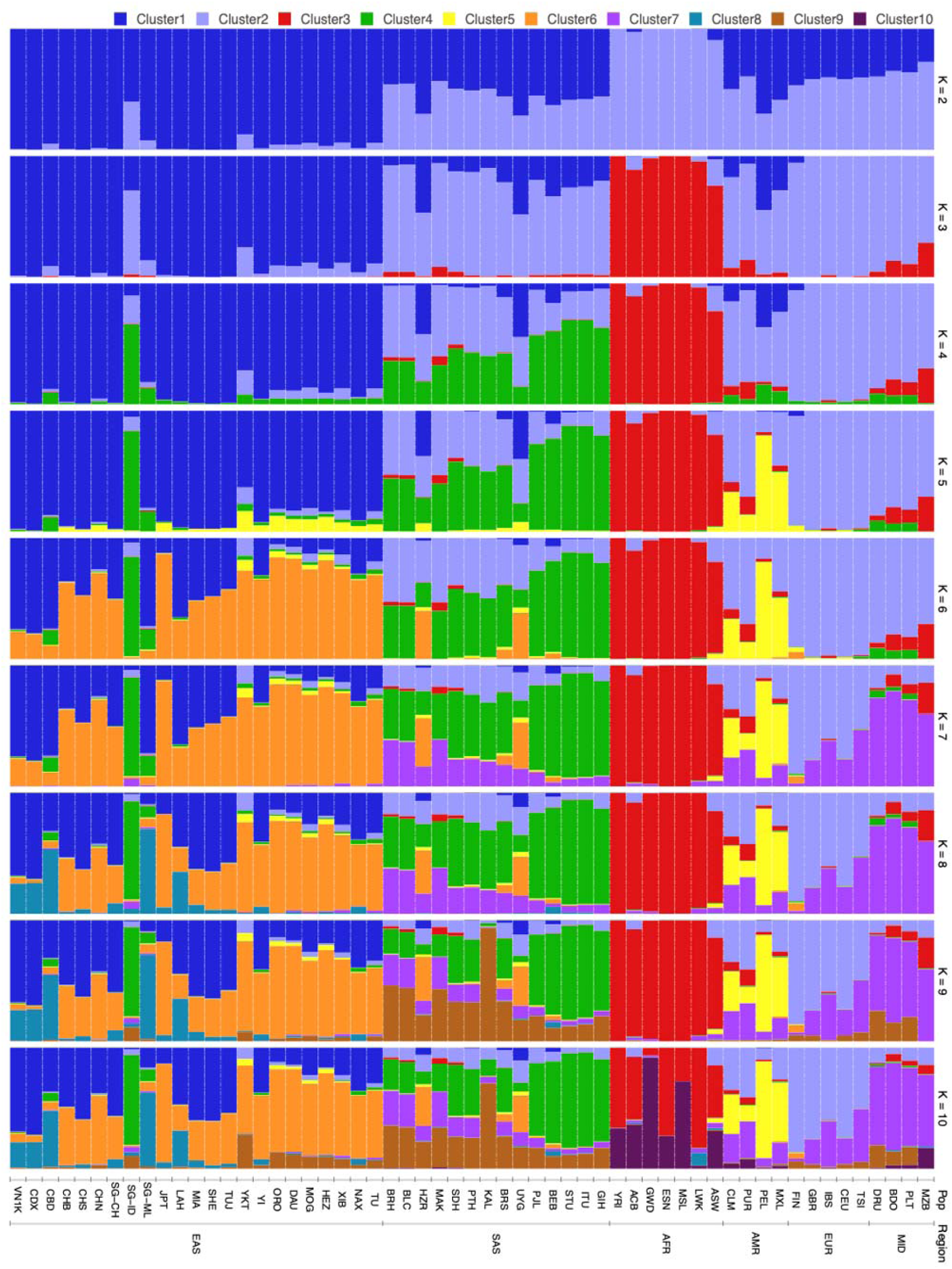
Vietnamese population analysis. The population structure signature of VN1K was shown based on the admixture ratio of ancestral components for each individual. The results highlighted the genetic diversity within the target population by comparing Fst values between Vietnam and other Asian groups in the HGDP project. VN1K (this dataset), CDX (Chinese Dai in Xishuangbanna), CBD (Cambodian), CHB (Han Chinese in Beijing), CHS (Southern Han Chinese), CHN (Chinese), SG-CH (Singapore Chinese), SG-ID (Singapore Indian), SG-ML (Singapore Malay), JPT (Japanese in Tokyo), LAH (Lahu), MIA (Miao), SHE (She), TUJ (Tujia), YKT (Yakut), YI (Yi), ORO (Oroqen), DAU (Daur), MOG (Mongolian), HEZ (Hezhen), XIB (Xibo), NAX (Naxi), TU (Tu), BRH (Brahui), BLC (Balochi), HZR (Hazara), MAK (Makrani), SDH (Sindhi), PTH (Pathan), KAL (Kalash), BRS (Burusho), UYG (Uyghur), PJL (Punjabi from Lahore), BEB (Bengali from Bangladesh), STU (Sri Lankan Tamil from the UK), ITU (Indian Telugu from the UK), GIH (Gujarati Indian from Houston), YRI (Yoruba in Ibadan), ACB (African Caribbean in Barbados), GWD (Gambian in Western Divisions), ESN (Esan in Nigeria), MSL (Mende in Sierra Leone), LWK (Luhya in Webuye, Kenya), ASW (African Ancestry in Southwest US), CLM (Colombian in Medellin), PUR (Puerto Rican in Puerto Rico), PEL (Peruvian in Lima), MXL (Mexican in Los Angeles), FIN (Finnish in Finland), GBR (British in England and Scotland), IBS (Iberian populations in Spain), CEU (Utah residents with Northern and Western European ancestry), TSI (Toscani in Italy), DRU (Druze), BDO (Bedouin), PLT (Palestinian), MZB (Mozabite).

### 8. VN1K data portal and multi-scale genome browser for integrated data resource

To enable the broad usage and impact of the current study, we have focused on making the VN1K data and results publicly available to the community through a comprehensive, user-friendly data portal named MASH (Management, Analysis, Sharing, and Harmonization) (http://genome.vinbigdata.org). MASH was designed as microservices with container-based infrastructure following the FAIR principle (Findable, Accessible, Interoperable, Reusable) to efficiently manage and analyze large and diverse data sources from different projects (**Figure 8**). MASH enabled searching data with gene names, rsID, variant ID, and other commonly genomic information. It also included a data harmonization module, displaying results for all common steps, including pre- and post-imputation quality controls, imputation, merging, and re-imputation. Importantly, MASH also integrated a genome browser which allowed researchers to work with both the linear (GRCh37/38) and graph genome reference (VGR), thus providing a dual architecture for visualizing and analyzing genomic variants and related information. This integrated system not only facilitated accessing data but also optimized performance of analytic tools for diverse communities for the purposes of research, education and clinical application.

**Figure 8.**
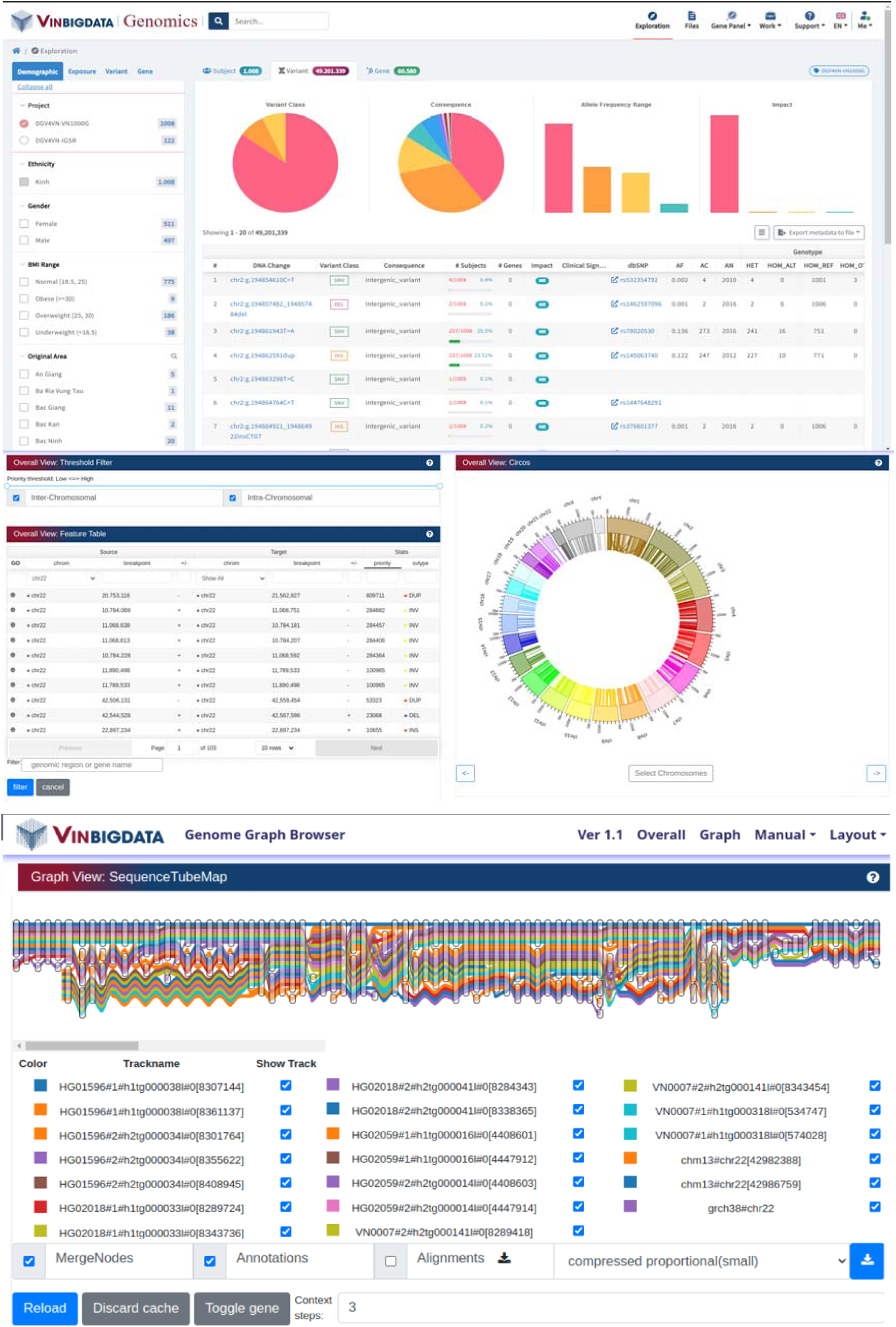
MASH data portal with web-based interface applications. (Top) The data portal, which facilitates researchers to explore individual- and population-level variants and genes, as well as filtering individuals (and their variants) by phenotypic traits. Users could also investigate various pre-built gene panels for HLA, PGx and common diseases as well as using various pre-built tools for data analysis and visualization. (Bottom) The integrated genome browser, which allows researchers to explore all SVs across chromosomes using both linear and graph-based reference. The variants could be filtered by genomic region or gene name and visualized using Circos software package. The multiple paths of SVs are shown by SequenceTubeMap as a graph genome reference.

## DISCUSSIONS

International human genome projects, such as the 1000 Genomes Project (1kGP) (1000 Genomes Project Consortium et al. 2015) and UK Biobank (Bycroft et al. 2018), have shed light on understandings of human genetic diversity and their association with different phenotypes and diseases. Such resources provided an estimate of variant allele frequency in different populations (MacArthur et al. 2012), which is critically important information to assess the likelihood and magnitude of causal variants of disease outcomes (Carlson et al. 2016) (Williams et al. 2018) (Santhiago 2016) and drug responses (Rueda-Clausen, Ogunleye, and Sharma 2015) (Kumar et al. 2014). These databases also facilitate the imputation of genotypes missing in genome-wide association studies (GWAS) and increase the statistical power to discover associations between diseases and low-frequency variants (Fadista et al. 2016) (Zhu et al. 2018). However, current studies have found that most genetic variants are rare and often occur at vastly different rates across populations, with many only appearing in specific populations (Hindorff et al. 2018) (1000 Genomes Project Consortium et al. 2015). In addition, as the majority of genomic studies to date have been focused on Caucasian ancestry (Sirugo, Williams, and Tishkoff 2019) (Mikhaylova and Thornton 2019) (Fatumo et al. 2022), millions of population-specific variants are missing. Therefore, genomic studies that capture local or specific population genetic makeups for specific racial and ethnic groups are essential.

Our study contributes to addressing the knowledge gap of human genetic diversity in specific racial/ethnic groups or populations, tackling the global issue in genomic equity and serving as robust genomic references not only for the 16th largest populations, but also for related populations of Asian-origin. With 1,011 individuals representing 54 of the total 63 administrative divisions of Vietnam, VN1K is the most up-to-date comprehensive database of genomic variants for the Vietnamese population. Although our study’s sample size is smaller compared to existing work in other countries, it remains the largest study for the Vietnamese population. We chose to study the Kinh ethnic group first for the VN1K project since they make up 86% of the entire population. Earlier studies have extensively explored the genetic architecture of Kinh people and other minorities using both single locus (Y-chromosome and mtDNA), genome-wide data and whole genomes (Macholdt et al. 2020) (Liu et al. 2020).

Our study provided the first graph-based genome reference named VGR for the Vietnamese population. VGR represents genetic diversity, while enhancing accuracy and sensitivity for detection of genomic variants, ranging from SNPs to large SVs. VGR incorporated all linear sequences from both short-read and long-read sequences of all participants, with the largest set of WGS, RNA sequencing, and long-read sequencing to date. Our analytics applied the latest methods to further capture the full genetic diversity of Vietnamese, including a deep learning approach and ensembles of results from a comprehensive collection of robust methods to build graph-based genomes. VGR helped visualize all paths of individuals and reduce reference bias of European ancestry from popular references.

Although the genetic variants have a substantial role in human evolution and population diversity, study of SVs in underrepresented populations is scarce. Using cutting-edge genomics sequencing technologies with PacBio and NanoPore, we were able to produce a comprehensive consensus reference for structural variants, a class of genetic variants that are increasingly recognized to impact hugely on the phenotypes, especially disease traits. Latest deep learning tools, shown to better identify SVs, were applied. In this study, we assessed all possible SV sets to achieve a consensus singular set including INS, DEL, DUP and ITX. Such results enabled us to construct the profiling of various disease-related SV sets and incorporate them into the pan genome reference.

VN1K provided functional analyses on various important regions of genome, in particular immunogenes, which has been long attracting great attention due to their critical roles in health conditions. Previous studies on the Vietnamese population have not used high quality, single-based resolution, whole genome analysis and were limited by small sample sizes. Using the high-depth whole genome sequencing data (30-60X), we identified a large number of novel HLA alleles (166 alleles) that were not identified in prior studies in Vietnam. Besides, the AF of most common alleles of both HLA-classes shown a high similarity with previous studies particular study performed in Kinh population (Vu-Trieu et al. 1997) (Hoa et al. 2008) (Que et al. 2022) (major ethnicity accounting for 86% country’s population [G.S.O.Vietnam]). Regarding KIR genetic variants, Amorim et al. (2018) recently reported an investigation of KIR distribution in 140 unrelated Vietnamese healthy people from Hanoi using standard PCR-SSP technology (Amorim et al. 2018), which is consistent with those observed from our study in the VN1K database, again suggesting the comprehensiveness of our database. The additional information gained by our study compared to the previous studies might be attributable to sample size, the sensitivity and the specificity of genotyping methods and bioinformatic tools **(Suppl. Table 4)**. The table presents the average allele frequencies as determined by the KPI software. Fisher’s exact test is employed to compare the allele frequencies across all variants of the KIR genes. The genes of 2DL1 and 2DS1 are different from VN1K to 1kGP-HC suggesting us to have further analysis on variants of these genes enabling advancing immunogenomics research, potentially leading to improved clinical implications, including treatment and management strategies for various disease outcomes.

In summary, we characterized the genomic variations in the Vietnamese population and provided the most comprehensive genomic database as the first pangenome reference for the Vietnamese population, to be used as a go-to resource for a wide-range of future genomic studies in Vietnam and beyond, especially in cross-ancestry analysis, taking into account populations of Asian-origin, consisting of 60% of the global population. This pioneering study for the Vietnamese population is expected to produce impacts extending beyond research to helping with education, providing potential social and healthcare benefits.

## METHODS

### Sample collection

We collected peripheral blood DNA samples of 1,050 healthy unrelated Kinh individuals (e.g., excluding donors known to be biologically related) from four provinces (Ha Noi, Da Nang, Ho Chi Minh city, Ha Tinh) which come from three main regions (North, Central, and South) in Vietnam. Kinh ethnic group was chosen because it is the largest ethnic group in Vietnam, accounting for 85.7% of the Vietnamese population (General Statistics Office of Vietnam’s 2023 report). The distribution or proportion of each group (cohort) is separated or classified based on the population growth data from individual city reports. This study was approved by the Institutional Board Review (or Ethics Committee) of the Vinmec International Hospital, Hanoi, Vietnam (543/2019/Q -VMEC).

The sample selection step was carried out using the health check examination provided by the Vinmec health-care service and the questionnaires. All participating individuals must be qualified by Vinmec physicians and sign a written consent form prior to joining the research. The metadata of samples with the de-identified samples were collected to use for research purposes only. Demographic information of participants was provided in the **Suppl. Table 1**. Quantitative traits were used to assess healthiness of individuals as provided in the **Suppl. Table 3**.

### DNA extraction, genotyping and sequencing

Genomic DNA was extracted and the quantification was performed using Qubit dsDNA Broader Range Assay Kit (Life Technologies) or Quantifluor dsDNA System (Promega). DNA amount > 1μg; DNA concentration > 35ng/μL. Library preparation was done using Illumina protocol TruSeq DNA PCR-free High Throughput Library Prep Kit with IDT for Illumina TruSeq DNA UD Indexes (e.g., UD index was used to reduce index-hopping). All library clean-up steps were performed following the standard protocol of the manufacturer (i.e., Illumina). Quantitative PCR (qPCR) was carried out using the KAPA Library Quantification Kits for Illumina platforms (Kapa Biosystems) before sequencing. Subsequently, intact genomic DNA was fragmented and sequenced at Hi-tech Center of Vinmec International Hospital. Paired-end 150bp whole genome sequencing with an insert size of 450 bp was performed on the Illumina Novaseq6000 platform. The target depth was 30x for all samples, with 100-120 GB sequencing data.

A subset of 96 samples from VN1K were randomly selected to be genotyped by The Axiom Precision Medicine Research Array (PMRA). Long-read sequencing was performed on another subset of our cohort on two platforms. PacBio Sequel II platform was conducted in two (02) reference samples (i.e., a female and a male sample) and four (04) pooled samples (i.e., 10 samples per pool) with 30X coverage. Next, we performed Oxford Nanopore Technology (ONT) sequencing for the aforementioned two reference samples with 30x coverage.

**Quality control (QC).** The raw fastq data files from each lane on Illumina Novaseq6000 platform were preprocessed by removing adapter sequences and trimming polyG in read tails which often happens in the Illumina two-color system. Furthermore, we used different filters on fastq files, including removing reads with over 50% bases having quality under Q30 or average base quality under Q30.

**SNP and INDEL calling.** The clean fastq files of each lane were then mapped against the reference genome GRCh38 p7 before being combined into one bam file. We used three state-of-the-art pipelines for variant calling: (1) Human_par pipeline which resembled bwa-0.7.12 and the GATK4 best practice pipeline with proper pseudo autosomal region ploidy values implemented on NVIDIA Parabricks (https://docs.nvidia.com/clara/parabricks/v3.5/text/human_par_pipeline.html), (2) Google DeepVariant on NVIDIA parabricks v3.5, (3) Illumina Dragen v3.7 that was analyzed locally on the Illumina Dragen On-Premise Server with a hardware-accelerated FPGA card. The reference genome GRCh38 p7 and the indexed files were downloaded from the resource bundle on the GATK website (https://console.cloud.google.com/storage/browser/genomics-public-data/resources/broad/hg38/v0/). To assure uniformity, we only kept the samples with the coverage at 15X that should be over 90% of the whole genome, and the improper pair percentage (IPP) be under 5%. We removed 79 samples that failed these filters. We further removed the other ten outliers outside the VN1K cluster, six duplicate samples, and two samples with contradictory sex declarations and inferred sex.

For each pipeline, GVCF files were combined to obtain a single cohort VCF files using explicit tools recommended by each development team: 1) GATK GenomicsDBImport to combine the Human_par GVCF files, 2) Google GLnexus to combine the DeepVariant GVCF files, and 3) Dragen Genotyper to combine Dragen GVCF files. Next, we applied GATK VQSR to the call set of GATK GenomicsDBImport and the default hard filter of Dragen and DeepVariant pipelines. Only passing sites of each pipeline were selected and further processed by taking the consensus of at least 2 pipelines.

**CNV calling.** Dragen CNV was used to call Copy Number Variants per each sample and then merged all the CNV, based on the reciprocal overlapping of at least 80%. All CNV calls from Dragen were filtered to pass all of four conditions, including 1) CNV with length below 10000, 2) CNV with quality below 10, 3) CNV with low supporting number of bins with respect to event length, and 4) CNV with copy ratio within ±0.2 of 1.0. The total CNVs passed all filters from 1011 samples is 628,301. The CNVs that are 80% or more overlapping were merged into one combined CNV. The start and end of combined calls are determined as the mean of start and end positions of individual CNVs. After merging, the combined list of CNVs consists of 20,241 CNV calls. To identify novel/replicated CNV, we compared with the structure variant of the Database of Genomic Variants released in 2020 (DGV) (MacDonald et al. 2014). We divided novel and replicated loci into three categories, including 1) Common > 5%, 2) Intermediate from 1% - 5%, and 3) Rare > 1%.

**Variant calling**: The resulting clean FASTQ files were then mapped against the reference genome GRCh38 p7 and combined into a single BAM file. For variant calling, we employed three state-of-the-art pipelines: 1) Human_par pipeline, resembling bwa-0.7.12 and the GATK4 best practice pipeline, implemented on NVIDIA Parabricks 1; 2) Google DeepVariant, also implemented on NVIDIA Parabricks v3.5 1, and 3) Illumina Dragen v3.7, analyzed locally on the Illumina Dragen On-Premise Server with a hardware-accelerated FPGA card. We used Dragen CNV to call Copy Number Variants (CNV) for each sample, merging them based on reciprocal overlapping of at least 80%.

We used a set of 96 PMRA genotyping data to evaluate the VN1K imputation reference panel. The remainder of the VN1K dataset, comprising 915 samples (named VN_915) was used as a reference panel. The imputation of combined panels utilized Whole Genome Sequencing (WGS) data from the 1000 Genomes Project with high coverage (1kGP-HC) and the Singaporean Genomes Project (SG10k) as a reference. The imputation pipeline was performed on chromosome 20 using Minimac4 with five imputation panels, including three individual panels (VN915, 1kGP-HC, SG10k) and two merged panels (VN915-1kGP-HC, VN915_SG10k). The genotyping data was then imputed compared to the WGS data to measure the R2, and NR-allele concordance rate.

In addition, long-read sequencing was performed on a subset of our cohort. First, we utilized PacBio Sequel II platform for two reference samples (a female and male samples) and 40 pooled samples (10 samples per pool) with 30X coverage. Secondly, we performed ONT sequencing for the aforementioned two reference samples with 30x coverage. These datasets not only enable better detection of SVs for the cohort but also provide DNA methylation information that could not be achieved with short-read sequencing data. SV calling was processed using pbsv 2.9.0 for PacBio data and CuteSV 2.0.2 <u>(Jiang et al. 2020)</u> for ONT data and then filtered by bcftools 1.13. In detail, we retained only the chromosomes within the range of [1-22XY], while discarding others. Subsequently, we employed the SURVIVOR (1.0.7) to merge the obtained SVs, setting a maximum measured pairwise between breakpoints limit of 1000 and length of SVs is at least 50 base pairs (bp).

**Structure variant calling**: We implemented a comprehensive genome assembly workflow that integrates long-read data from PacBio and ONT platforms, as well as short-read data from Illumina platforms, using a hybrid assembly approach with the Wengan software (Di Genova et al. 2021). The resulting contigs had quality assessment with QUAST (Gurevich et al. 2013) and were then further assembled into scaffolds using the GRCh38 reference genome as a guide, facilitated by the ragtag tool. Any remaining gaps in the scaffolds were filled using the hg38 reference, producing the final genome assembly. These assembly references were then used to progressively add to a pan-genome graph, in order to incorporate the specific genomic value of the Vietnamese population. By aligning the Vietnamese genome data against our new reference, we could identify novel sequences and SVs and seamlessly integrate them into the expanding pan-genome graph. The deep learning workflow includes four steps: First, quality control is performed for all structural variant (SV) data, such as PacBio and Nanopore. The criteria involve filtering, read depth, length, and quality score of SV. Second, a deep learning model named cnnLSV is used to convert each SV into an image and predict SV types. The training process for the cnnLSV model includes two stages: filtering and detecting. In the filtering stage, existing callers were used to refine detection results, labeling true and false positive SVs, and converting them to images. Next stage, the images were applied to train models to detect structural variants. Finally, a consensus SV dataset is created by merging all SV positions using SURVIVOR. This workflow ensures the quality of the SV by incorporating an image-based approach for better detection of SVs in VN1K.

**Pangenome graph construction:** To build the pangenome graph, we first implemented a comprehensive genome assembly workflow that integrated long-read data from PacBio and ONT as well as short-read data from Illumina platforms, using a hybrid assembly approach with the Wengan tool. The resulting contigs had quality assessment with *QUAST* were then further assembled into scaffolds guided by GRCh38, facilitated by the *ragtag* tool, to fill the remaining gaps and produce the final genome assembly. Then, the Vietnamese Pangenome Graph Reference (VGR) comprises core assemblies and incorporates variants derived from both long-read and short-read sequencing data of VN1K. For the core pangenome assemblies, the *PanGenome Graph Builder (PGGB) v0.6.0* was used to create a graphical fragment assembly (GFA). This tool processed multiple genome assemblies as input, including the GRCh38, the recently released CHM3-T2T assembly, two KHV assemblies, and two VN1K assemblies produced using a hybrid approach combining PacBio and ONT technologies. Subsequently, *vg v1.156* was employed to integrate the identified genetic variations (SNPs, INDELs, and SVs) identified from 1008 VN1K and four pooled long-read PacBio datasets.

**Methylation analysis**: A DNA methylation reference panel was constructed using data from two healthy Vietnamese samples (i.e., one male and one female) based on the best long-read sequencing technologies from ONT and PacBio. ONT data was generated from R10.4.1 chemistry which outperforms R.9.4.1 in methylation calling, while PacBio data utilized HiFi reads. Each technology followed the respective manufacturer’s best practices for DNA methylation calling, as detailed in **Suppl. Methods**. In the final step, methylation scores were called from 5mC BAM files based on the MM and ML tags. To standardize the final format of the two best practices, we used Modkit to generate the final BEDGraph files for both technologies. Subsequently, for each sex, we created a consensus by overlapping sites with common positions based on chromosome, start, and end coordinates. The final DNA methylation reference panel was constructed by overlapping the consensus files of the two sexes, with chrY taken directly from the male consensus file. Next, we compared the Vietnam methylation reference to other populations. Due to the differences in technologies between the VN1K and 1kGP methylation datasets, we corrected batch effects using the *limma* package. To identify differentially methylated regions (DMRs) between Vietnamese and other populations, methylation sites were categorized as hypermethylated and hypomethylated based on their methylation levels (>0.75 for hypermethylation, <0.25 for hypomethylation). Sites that were consistently hypermethylated in the Vietnamese population but hypomethylated in at least one other population of African (AFR), Admixed American (AMR), East Asian (EAS), European (EUR), South Asian (SAS) or vice versa were retained for further analysis. Enrichment analysis was then performed using Fisher’s exact test in the LOLA package, and gene annotation was carried out with the *annotatr* package to explore patterns within the DMRs. Finally, the gene expression levels of these DMR-associated genes were validated through RNA sequencing data from 14 control samples, comparing them to the gene expression profiles of corresponding population samples from the 1kGP dataset.

**Immunogenomic and pharmacogenomic analysis:** Jost’D distance was used to calculate the distance between VN1K and other populations based on pharmacogenes. WGS-derived HLA sequence data of each individual in VN1K and 1kGP-HC were imported in xHLA for both HLA class I and class II genotyping with resolution of 4 digits, whereas the genotype of KIR complex was called by two bioinformatic tools, KPI <u>(Roe and Kuang 2020)</u> and KIRCLE.

The equation for Jost’s D is:

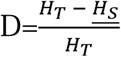

where:

- H_T_ is the total heterozygosity (genetic diversity) in the combined population.
- H_S_ is the average heterozygosity within subpopulations.

**GWAS analysis:** A genome-wide association study was performed with 810 samples with a full lab test for 12 numerical traits. Under an additive genetic model, GWA analysis was carried out using linear regression using best practices (e.g., linear regression is used to test the association between genetic variants and quantitative traits which were collected in the study **(Suppl. Table 3)**. Covariates considered were year of birth, gender, weight, height and the first 10 principal components. PCA were then performed from post-processed variants that were pruned via PLINK v1.9. The results were further filtered with the threshold of 5.86*10^9. The final set of loci were investigated with VEP Ensembl and GWAS catalog (ver. 2022-12-21).

**Reference panel construction:** We have constructed a haplotype reference panel for phasing and imputation based on the database of genetic variants created by the VN1K project. From the final dataset, the singleton variants were removed using VCFtools version 0.1.16 (Danecek et al. 2011). We filtered out multi-allelic sites via the BCFtools norm tools version 1.9 (H. Li et al. 2009). SHAPEIT4 (Delaneau et al. 2019) was used to phase the haplotype of all samples with default settings. Finally, the result was formatted with Minimac3 (Das et al. 2016) for imputation.

The 1kGP reference panel (High coverage version) was downloaded from sites (https://www.internationalgenome.org/) and SG10K was from Precision Health Research, Singapore (https://www.npm.sg/collaborate/partners/sg10k/). They were processed the same as the VN1K reference panel. Two merged panels were respectively merged from our VN1K dataset with the 1kGP and SG10K following guidance from a study by Huang et al. (Huang et al. 2015). In carrier screening, we found genes were not featured in the ACMG prenatal screening for genetic disorders (e.g., pathogenic variants of *PMFBP1*, *MTSS2*, *C9, SRD5A2*, and *HFE)* with considerable AF in the study (**Suppl. Figure 4**) which have important implications in genetic screening assays for the Vietnamese population.

**Pre-phasing and imputation:** Pre-phasing was performed by SHAPEIT4 (Delaneau et al. 2019) with the default setting. Minimac4 (Das et al. 2016) was used to impute phased variants with our haplotype reference panels. Reference panel evaluation for imputation. In order to analyze and compare, we used five imputation panels, including 1kGP, the SG10K, our own VN1K panel, and two combination panels that combined the VN1K datasets with the 1kGP and the SG10K, respectively. 96 unrelated Vietnamese individuals were randomly selected and genotyped with Axiom Precision Medicine Research Array (PMRA) (Thermo Fisher Scientific, MA, USA) for evaluation purposes. Imputation accuracy was measured with aggregate R2 (Pearson correlation) between the dosage of imputed genotypes and the true genotype of variants which was sequencing data. We computed the aggregated R2 in each MAF group for comparison purposes specifically for rare and low-frequency variations, which are frequently challenging to impute effectively.

**HLA imputation panel construction:** Inference of HLA allele from SNP data is a non-trivial task. Many SNPs with high linkage disequilibrium are likely to be presented together. Besides, the number of existing HLA alleles is tremendous, leading to multiple sequences with similar subsequences. These phenomena interfere with the input signal and create non-independent observations; thus, the analytical model is required to be able to filter out the noise, keeping only the important signals. In SNP microarray data, the problem is further elevated, because only some representative variants are presented on the chip. We develop a processing model combined with a principal component analysis (PCA) algorithm for input data with the aim of retaining the SNPs that carry the most valuable information and reduce the input dimensions at the same time. The problem of linkage disequilibrium is handled through the ability of convolutional neural networks, which represent data information with different layers, and efficiently aggregate information from neighbor SNPs through the sliding window. Based on biological knowledge, we built a separate model for each group of HLA genes according to their position on the gene. The HLA alleles were called by using 3 different methods and consented through majority voting (Xie et al. 2017) (Lee and Kingsford 2018) (Kim et al. 2019). In the data preparation process, although it is possible to identify the haplotype of the HLA gene and the genetic variant, it is not possible to determine if the exact orders of these two haplotypes are similar. In order to preserve diplotype information and ignore position information, we encode them with 2 operations, AND and OR. The AND operation indicates whether both haploids have variation at this location while the OR operation indicates whether one variation on both haploids. With this encoding, we eliminated the workload of generalizing HLA information from the exact haplotypes and made the model work on the convolutional network most effectively.

The HLA and variant data from VN1K are randomly divided with a ratio of 9-1 to obtain the training and validation dataset. The validation dataset will be used to adjust the model’s hyperparameters, and the final hyperparameters for our model are specified in the **Suppl. Methods.** Finally, we use the Binary Cross Entropy (BCE) loss function.

**Population genetic structure analyses:** We used the liftOver (Genovese et al. 2024) program to convert the coordinates of the SNPs from hg37 to hg38 (in the SG10K). After this, there were reference alleles in the SG10K that must be changed to alternate alleles in hg38 coordinates, and vice versa. Finally, the SG10K dataset included 9,254 individuals belonging to 108 populations and 1,627,296 SNPs.

To merge all SNPs from the VN1K, 1kGP, HGDP and SG10K dataset, we used Bcftools (H. Li et al. 2009) program to extract all variants called in four datasets then merged the VCF file.

After PCA analysis from VN1K and 1kGP, KHV samples from 1kGP seemed close to VN1K, we decided to exclude 100 KHV samples in the 1kGP to mainly focus on our dataset. In addition, because the large number of samples in SG10K was overrepresented for just 3 ethnicities, we randomly selected 200 samples for each population in this project for our analysis (Willing, Dreyer, and van Oosterhout 2012) (**Suppl. Figure 8**). We also removed some populations in the HGDP dataset that had under five samples. The final dataset had 4626 individuals belonging to 57 populations.

Before ADMIXTURE analysis on 57 populations, we filtered out SNPs with MAF ≤ 0.05 or the missing rate ≥ 0.05 then pruned SNPs with high correlations using indep-pairwise in PLINK. The final genotypes of 57 populations included independent 130120 SNPs in PLINK binary format.

Then, at each K from 2 to 35, the files were analyzed by the ADMIXTURE 10 different seed values with 5-fold cross-validation. We chose the best optimal cross-validation error at each K and took an average of each cluster in each population after using CLUMPAK (Kopelman et al. 2015) program to analyze results from different seeds. Finally, we visualized the posteriors by the pophelper library (Francis 2017) in R.

We performed PCA analysis by PLINK (Purcell et al. 2007) on VN1K dataset after pruned variants with high linkage disequilibrium and MAF ≤ 0.01. We split our samples by their gender and original region to visualize different principal components.

**VN1K data portal construction:** Leveraging MoMI-G (Yokoyama et al. 2019) as the underlying framework, the Genome Graph Browser offered a user-friendly interface to visualize SVs by a variation graph. The browser boasted a rich set of functionalities, including functionalities to explore SV distribution across chromosome, path graph, and gene annotation. To achieve these visualizations, SVs were called to compute into a dedicated dataset. This dataset was then displayed on a variant table tab and visualized using a Circos plot. Furthermore, SequenceTubeMap was assembled into the genome graph browser for SV graph visualization. Gene annotation function was designed to retrieve from Ensembl or regions listed in BigBed format of the hg38 human reference genome. Gene annotation information was shown downloading as a BED file. For input, the browser accepted variation graphs in XG format, along with the metadata associated with the SVs. Additionally, it could be processing graph alignment information (GAM) derived from BAM files. All these inputs were conveniently specified within a configuration file using YAML format. In summary, the user interface was built using TypeScript and implemented for testing on Linux operating systems. Finally, a data portal called MASH (Management, Analysis, Sharing, and Harmonization) was developed with optimized modules for storing and sharing data, and providing a user-interface to explore all variant information in VN1K. MASH also included a genome browser which integrated the reference genomes, both linear-based references GRCh37/GRCh38 and graph-based references VGR.

## Data availability

All the data and results could be found in the Main Text, Supplementary Files, and the MASH Data Portal (genome.vinbigdata.org). Please contact the corresponding authors for further request.

## Author Contributions

Initials of authors in the list (the same order): TTHT, THH, MHT, NTN, DTN, TMP, NNN, GMV, VCD, QTV, TKN, SVN, HQV, TMN, TD, HN, TD, CL, HTTN, NQL, QHN, LTL, TP, MD, DMV, HTTL, TDN, LTN, YH, DXD, GHP, TT, LN, CTH, HNL, VSL, LL, NTD, DHL, QN, VHV, NSV.

NSV, QN, DHL, NTD, LL, and VSL conceptualized, designed, and supervised the project. VHV, MD, TT, and LTN supervised the project. TTHT and THH conducted the primary data analysis and interpretation. NTN, DTN, TMP, NNN, GMV, VCD, QTV, TKN, SVN, HQV, TMN, HTTN, NQL, QHN, and CTH contributed to the data analysis. MHT, TP, LTL, DXD, and GHP performed the data curation. MHT, DMV, HTTL, TDN, LTN, YH, DXD, GHP, and TT contributed to the sample collection. DMV, HTTL, and LN conducted the sequencing experiments. TD, HN, TD, and CL built the MASH Data Portal. TTHT, THH, NTN, DTN TMP, NNN, QTV, GMV, VCD, TKN, SVN, QN, and NSV drafted the manuscript. HNL, VSL, NTD, and DHL revised the manuscript. NSV and QN revised, rewrote, and wrapped the final manuscript. All authors read and approved the final manuscript.

## Acknowledgements

We would like to thank Drs. Nong Van Hai and Nguyen Dang Ton (Institute of Genome Research, Vietnam Academy of Science and Technology) for their valuable comments on data analysis. We are especially grateful to Drs. Chi V. Dang (Ludwig Institute for Cancer Research, Johns Hopkins University School of Medicine), Michael Beer (Department of Biomedical Engineering and Department of Genetic Medicine, Johns Hopkins University), Ken Chen (Department of Bioinformatics and Computational Biology, The University of Texas MD Anderson Cancer Center), Pavel Pevzner (Department of Computer Science & Engineering, University of California, San Diego), and George P. Patrinos (Department of Pharmacy, University of Patras) for their insightful comments on the final manuscript. We also thank Dr. Quang Tran (Department of Bioinformatics and Computational Biology, The University of Texas MD Anderson Cancer Center) and the intern students at Vingroup Big Data Institute and GeneStory who contributed to the data analysis and wetlab experiments. Our sincere appreciation goes to all the individuals and their families who generously agreed to participate in this research. We are grateful to the Vinmec Healthcare System and Hanoi Medical University Hospital for their support with sample collection, sequencing facilities, and clinical expertise. This project was primarily funded by Vingroup Big Data Institute and partially supported by VINIF Grant VINIF.DA.2020.02.

