## Supplementary Methods for "VN1K: a genome graph-based and function-driven multi-omics and phenomics resource for the Vietnamese population"

*This file includes Supplementary Methods 1-5, which give more details for Methods session.*

#### 1. Workflow for variant calling from long-read sequencing

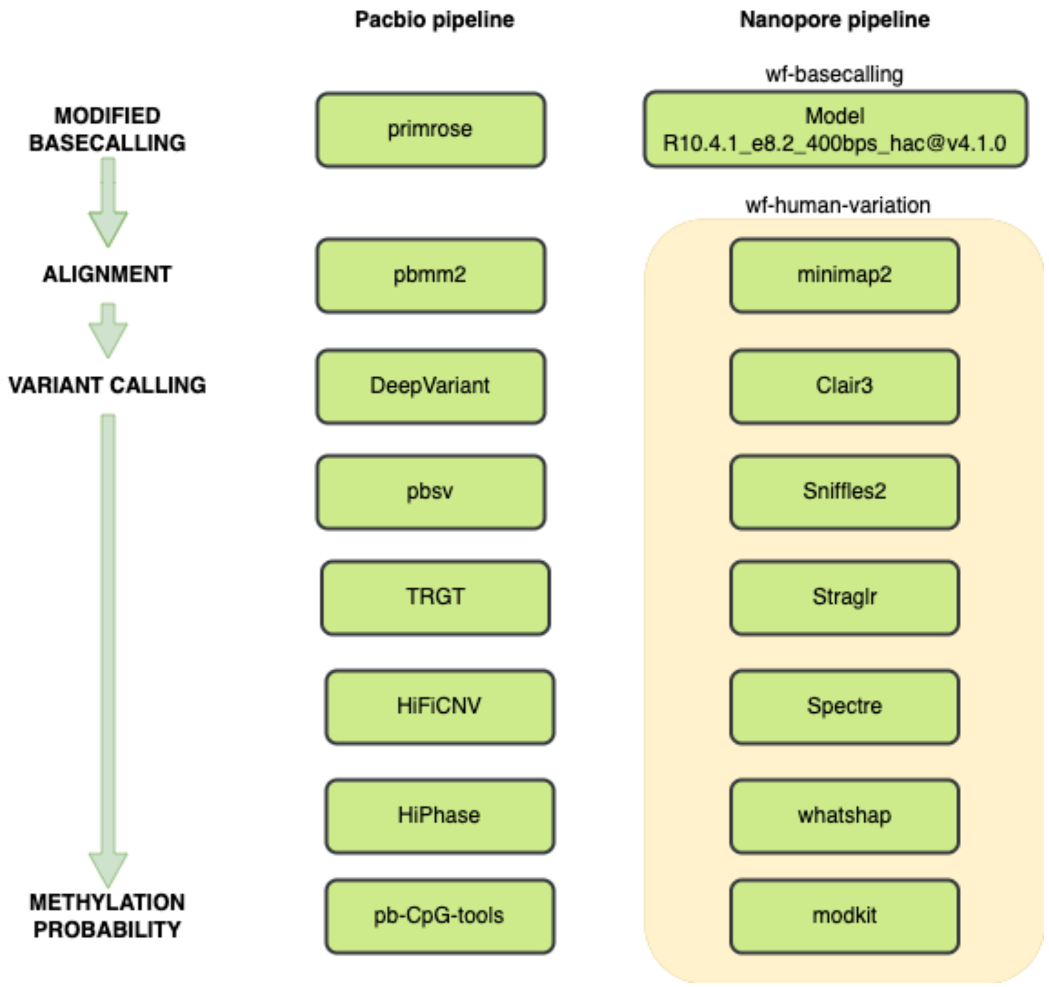


**Figure 1.** The diagram illustrates the comprehensive workflow starting from modified base calling to alignment, variant calling, and methylation probability estimation.

### 2. Deep learning framework

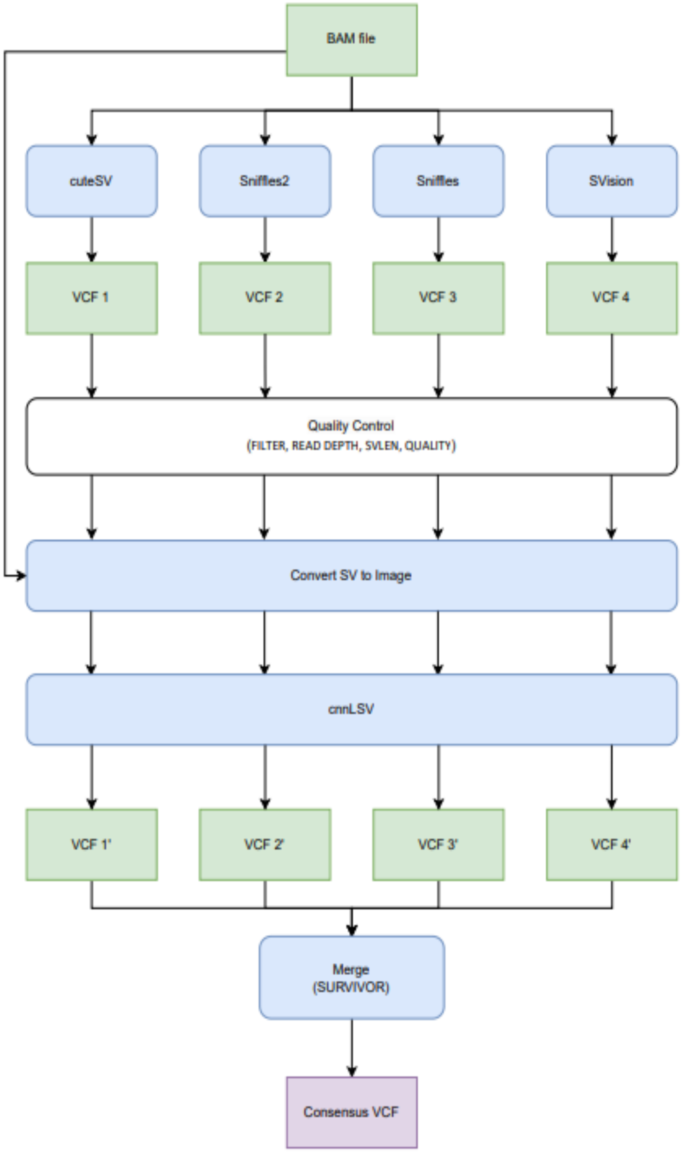


**Figure 2**: The diagram shows the workflow for obtaining a consensus set of SVs by using deep learning and ensemble multiple tools.

The framework to detect SVs on long-read datasets is illustrated in **Figure 2**. Its pipeline was designed with three main phases. Phase 1 called SV from the BAM file and exported to VCF using different tools such as cuteSV, SVision, and sniffles. Next, a quality control step was applied to

VCF files, filtering out low-quality SV based on read depth, SVLEN, and quality of variants. Each SV in filtered VCF and BAM file as a reference panel was then converted to an image and fed into a deep learning model (cnnLSV). The model predictions were combined by using SURVIVOR to consolidate overlapping or conflicting calls and obtain a consensus set of SVs.

The framework is to detect SVs on long-read datasets. The long-read data for INS, DEL, INV, and DUP variants was encoded into images. A trained model then evaluated these images to identify true positives. If the model's final output value exceeds 0.5, the image is classified as a true positive; otherwise, it is considered a false positive. In this context, a true positive indicates the presence of the variant in long reads. We keep the true positives and discard the false positives for INS, DEL, INV, and DUP. Unlike these four variants, the variant call format (VCF) file generated by callers records TRAs at the breakpoint level, making it challenging to encode TRAs occurring between two chromosomes into images.

#### 3. Analysis of immunogenomic variants

HLA: vWGS-derived HLA sequence data of each individual in VN1K and 1kGP-HC were imported in xHLA for both HLA class I and class II genotyping with resolution of four digits. Fisher's exact test was used to assess the differences in allele frequencies between Vietnam (VN1K) and global populations (from 1kGP-HC). Fisher exact test $p < 0.05$ is the threshold for a significant difference.

KIR: Due to the complexity of variations, including gene content (haplotype) variability and allelic diversity, the genotype of KIR complex was called by two bioinformatic tools, KPI and KIRCLE. KPI systematically investigated gene-specific small markers (k-mer) of KIR genes and then applied those markers to a synthetic KIR probe algorithm to identify the presence/absence of 16 KIR genes and 16 haplotype structures. KIRCLE, on the other hand, was used to infer KIR alleles and genotype predictions for each KIR gene from aligned WES data in the form of a BAM or CRAM file. KIRCLE was performed through four major steps: 1) pre-processing, 2) local alignment with BLAST, 3) bootstrapped expectation-maximization, and 4) thresholding.

#### 4. Reference panel for phasing and imputation

We have constructed a haplotype reference panel for phasing and imputation based on the database of genetic variants created by the VN1K project. From the final dataset, the singleton variants were removed using VCFtools version 0.1.16. We filtered out multi-allelic sites via the BCFtools norm tools version 1.9. SHAPEIT4 was used to phase the haplotype of all samples with default settings. Finally, the result was formatted with Minimac3 for imputation.

The 1KGP reference panel (High coverage version) was downloaded from sites (https://www.internationalgenome.org) and SG10K was from Precision Health Research, Singapore (https://www.npm.sg/collaborate/partners/sg10k). They were processed the same as the VN1K reference panel. Two merged panels were respectively merged from our VN1K dataset with the 1KGP and SG10K following guidance from a study by Huang et al.

#### 5. Reference panel evaluation for imputation in the Vietnamese population

In order to analyze and compare to existing panels, we used five imputation panels, including 1KGP, the SG10K, our own VN1K panel, and two combination panels that combined the VN1K datasets with the 1KGP and the SG10K, respectively. 96 unrelated Vietnamese individuals were randomly selected and genotyped with Axiom Precision Medicine Research Array (PMRA) (Thermo Fisher Scientific) for evaluation purposes. Imputation accuracy was measured with aggregate R2 (Pearson correlation) between the dosage of imputed genotypes and the true genotype of variants which was sequencing data. The number of high-quality variants is measured with Rsq value. The Rsq metric measured how well the variants were imputed from the reference panel (assuming Hardy-Weinberg equilibrium). We counted the number of variants having Rsq≥0.8 (high-quality variants) in each MAF group for comparison purposes specifically for rare and low-frequency variations, which are frequently challenging to impute effectively. We evaluate the imputation error rate on 24 Asian populations from the Human genome diversity project (HGDP) data which contains the genotype from a microarray named "Illumina HuHap 650k". Fst statistics is an estimate calculated in accordance with Weir and Cockerham's 1984 paper via VCFtools. All computations were performed on chromosome 20.

**6. HLA imputation panel construction**

Many SNPs with high linkage disequilibrium are likely to be presented together. Besides, the number of existing HLA alleles is large, leading to multiple sequences with similar subsequences. These phenomena interfere with the input signal and create non-independent observations; thus, the analytical model is required to be able to filter out the noise, keeping only the important signals. In SNP microarray data, the problem is further elevated, because only some representative variants are presented on the chip.

We developed a processing model combined with a principal component analysis (PCA) algorithm for input data with the aim of retaining the SNPs that carry the most valuable information and reduce the input dimensions at the same time. The problem of linkage disequilibrium was handled through the ability of convolutional neural networks, which represented data information with different layers, and efficiently aggregate information from neighbor SNPs through the sliding window. Based on biological knowledge, we built a separate model for each group of HLA genes according to their position on the gene. The HLA alleles were called by using 3 different methods and consented through majority voting. In the data preparation process, although it was possible to identify the haplotype of the HLA gene and the genetic variant, it was not possible to determine if the exact orders of these two haplotypes were similar. To preserve diplotype information and ignore position information, we encoded them with two operations, AND (and) and OR (or). The AND operation indicated whether both haploids have variation at this location while the OR operation indicates whether one variation on both haploids. With this encoding, we eliminated the workload of generalizing HLA information from the exact haplotypes and made the model work on the convolutional network most effectively.
