## Supplementary Figures for "VN1K: a genome graph-based and function-driven multi-omics and phenomics resource for the Vietnamese population"


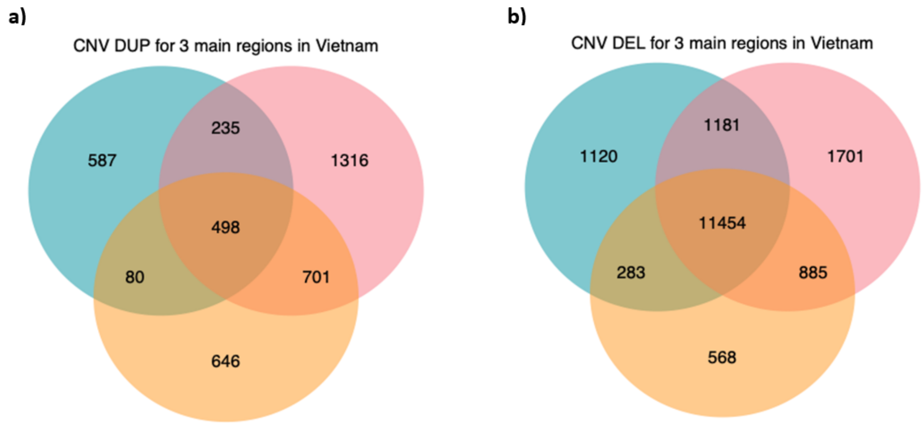


**Supplementary Figure 1.** The number of DUP and DEL CNV in three main regions of Vietnam.

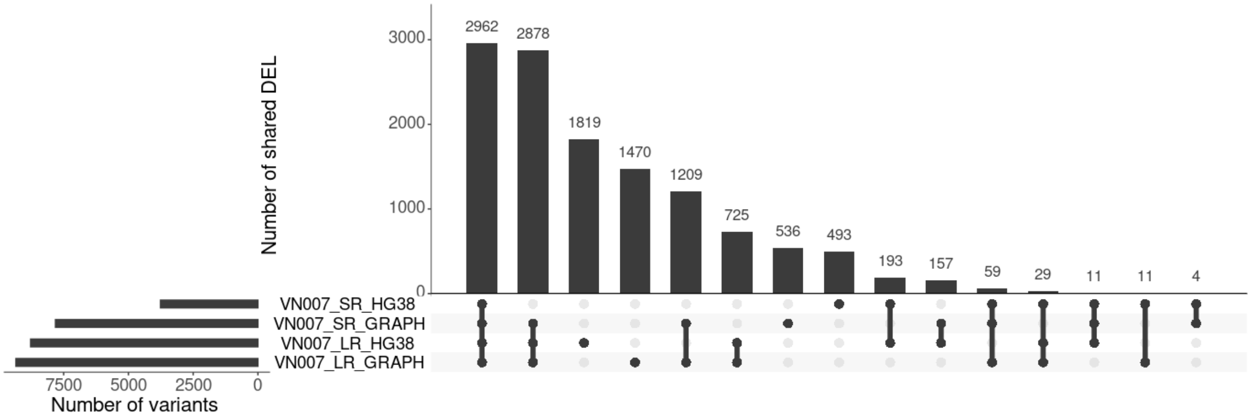


**Supplementary Figure 2.** Evaluation of SV calling by graph-based reference (VGR) and linear-based reference (GRCh38) with sample VN007. SR - short read sequencing, LR - long read sequencing, HG38 - GRCh38 linear reference.

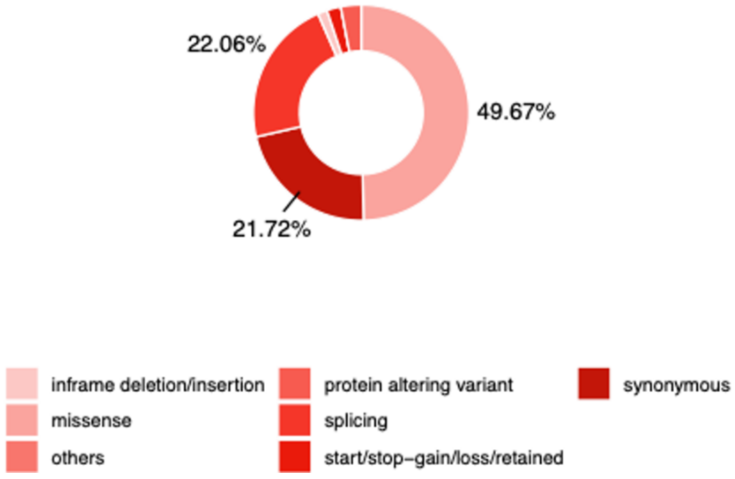


**Supplementary Figure 3.** Summary of the variant consequence of the VN1K's novel set.

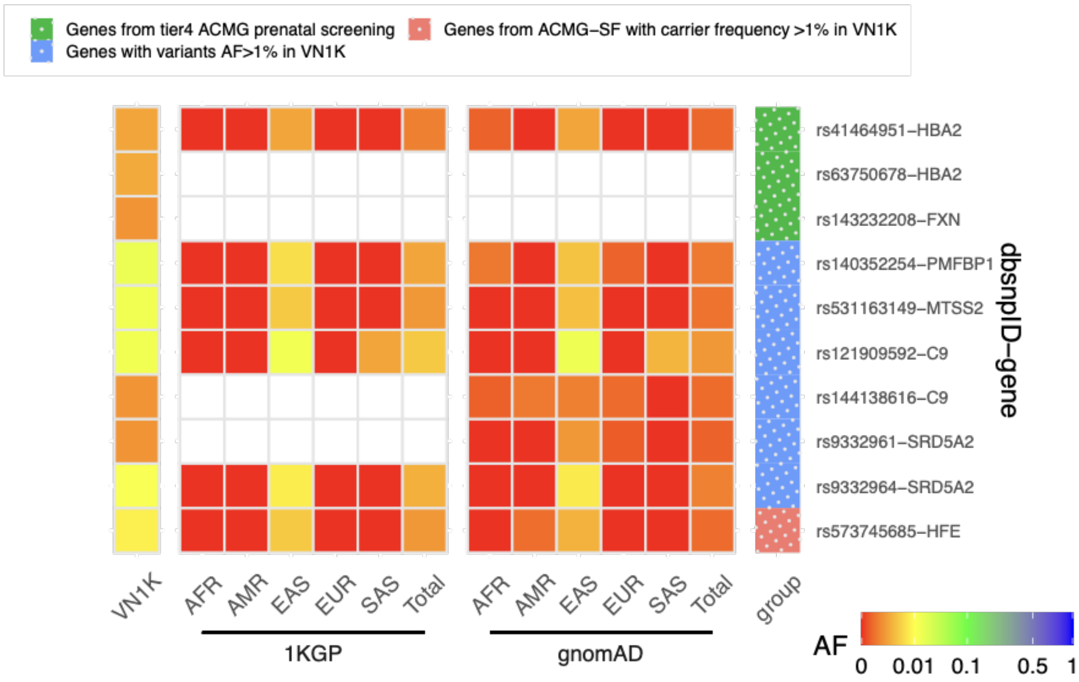


**Supplementary Figure 4.** Five genes (PMFBP1, MTSS2, C9, SRD5A2 and HFE) that do not appear in ACMG prenatal screening for genetic disorder with high allele frequency in VN1K.

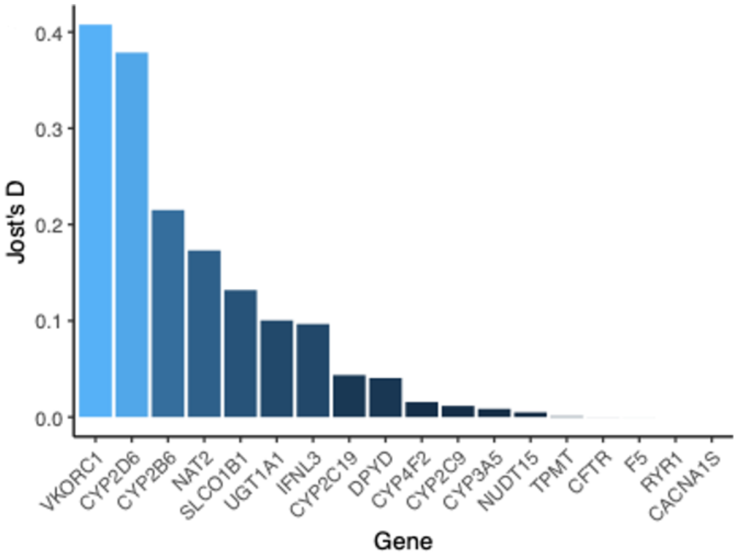


**Supplementary Figure 5**. Top pharmacogenomics genes with highest Jost's D genetic distance between VN1K and 1kGP-HC.

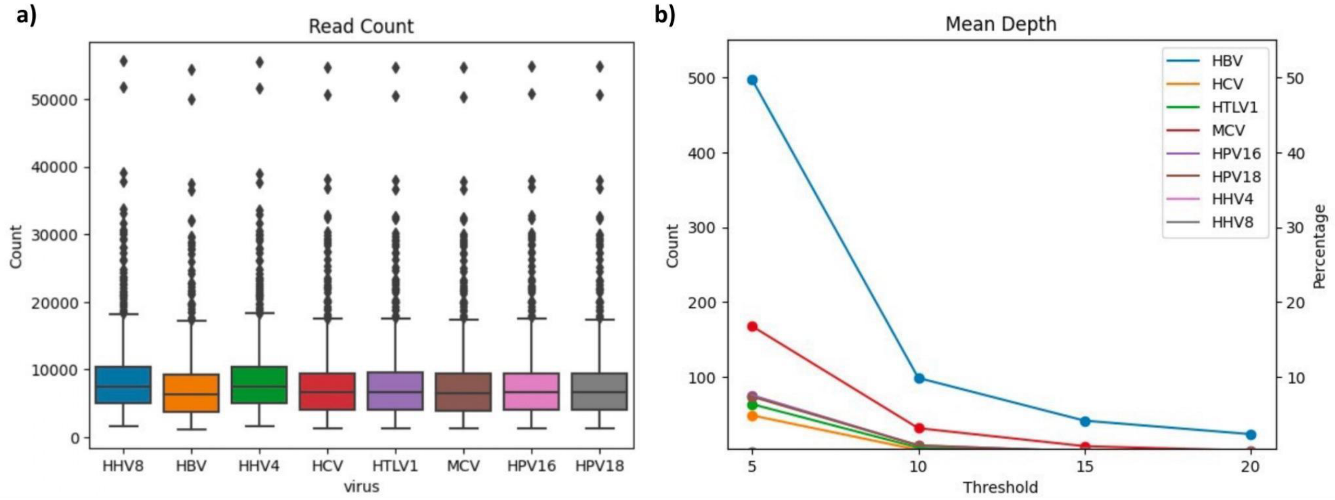


**Supplementary Figure 6.** The number of reads mapping with virus reference genomes. a) The number of VN1K human genome reads mapping with 8 virus genomes. b) The number of VN1K samples have mapped reads with different mean depth thresholds.

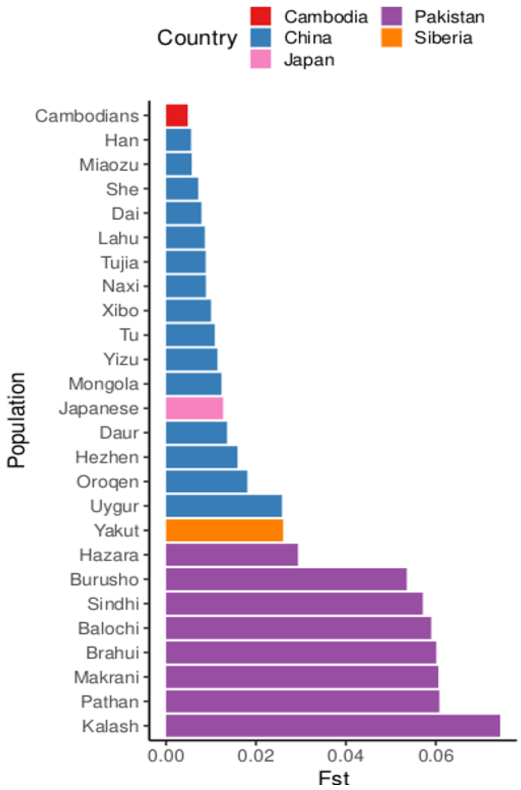


**Supplementary Figure 7**. Fst between Vietnam and other Asia group in HGDP project.

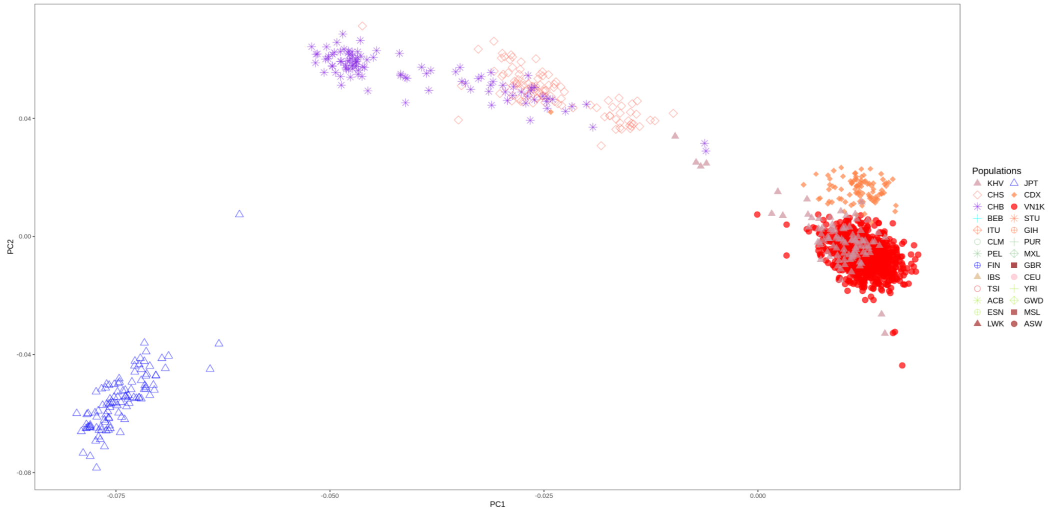


**Supplementary Figure 8.** PCA results VN1K and 1KGP samples focused on East Asia. . KHV: Kinh Vietnamese, JPT: Japanese, CHS: Southern Han Chinese, CDX: Chinese Dai, CHB: Han Chinese, VN1K: This dataset, BEB: Bengali, STU: Sri Lankan, ITU: Indian, GIH: Gujarati, CLM: Colombian, PUR: Puerto Rican, PEL: Peruvian, MXL: Mexican-American, FIN: Finnish, GBR: British, IBS: Spanish, CEU: Utah residents with Northern and Western European ancestry, TSI: Tuscan, YRI: Yoruba, ACB: African-Caribbean, GWD: Gambian, ESN: Esan, MSL: Mende, LWK: Luhya, ASW: African-American SW.

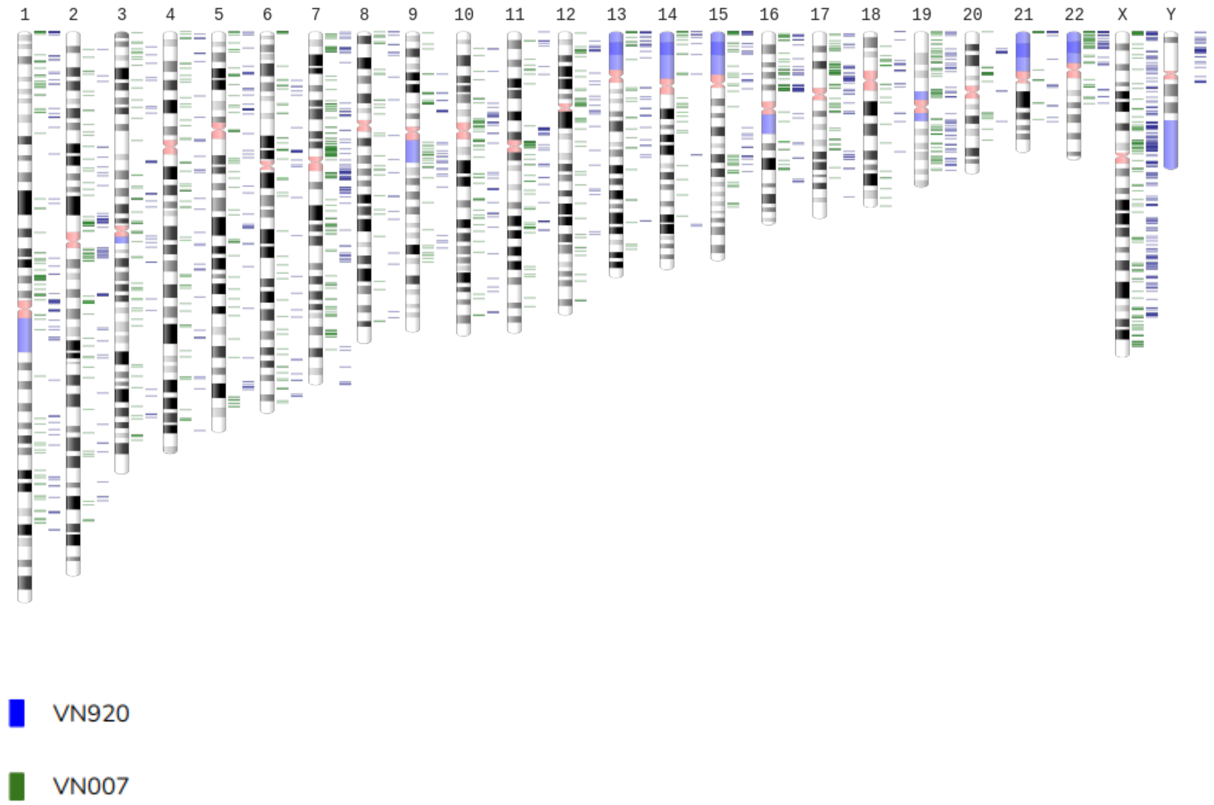


**Supplementary Figure 9.** The bands (G bands) correlate approximately with the DNA sequence underlying it: AT-rich areas stain darkly, GC-rich areas lightly. An idealized picture of such a G-banded karyotype (called an ideogram). About 400 dark bands per haploid genome are seen in this way. Two samples VN920 and VN007 are shown in the figure.

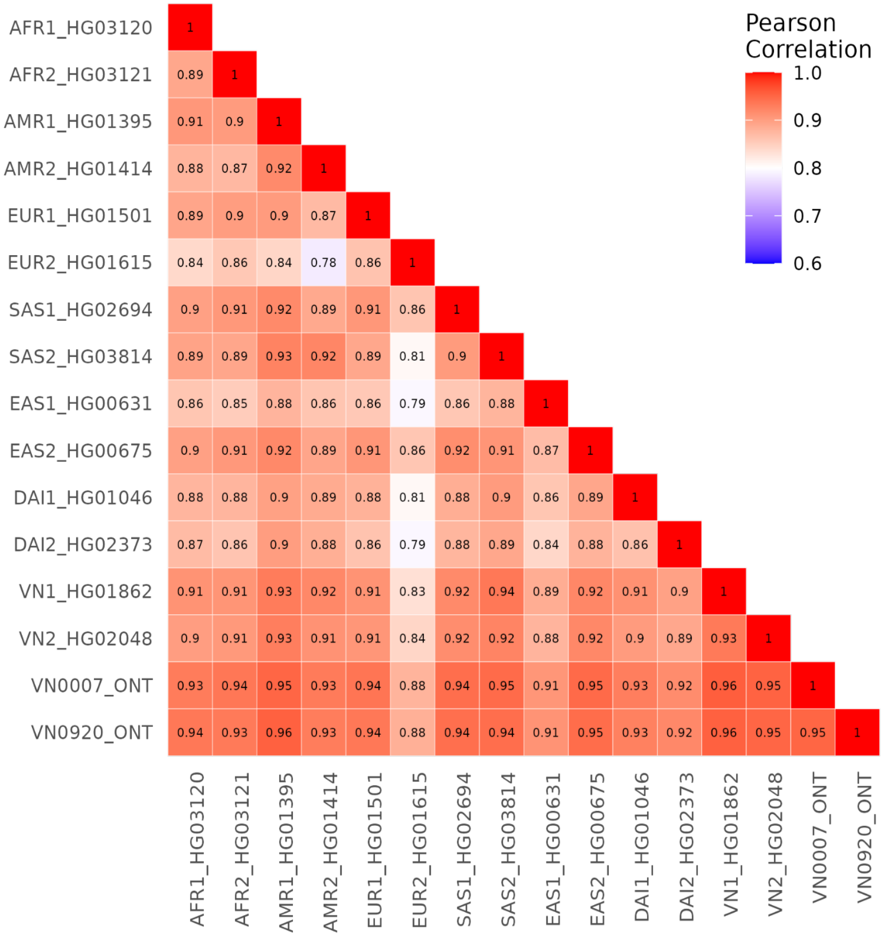


**Supplementary Figure 10.** Pearson correlation between 2 Vietnamese VN1K samples vs 1KGP samples after Batch Correction.

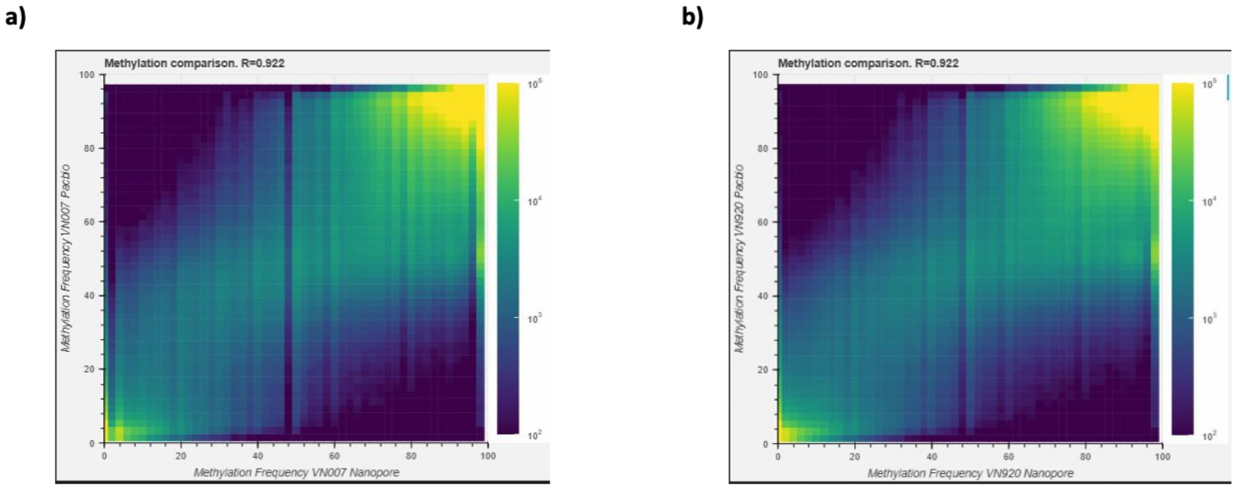


**Supplementary Figure 11.** Comparison of Methylation Frequencies between Pacbio and ONT for two Vietnamese Samples.

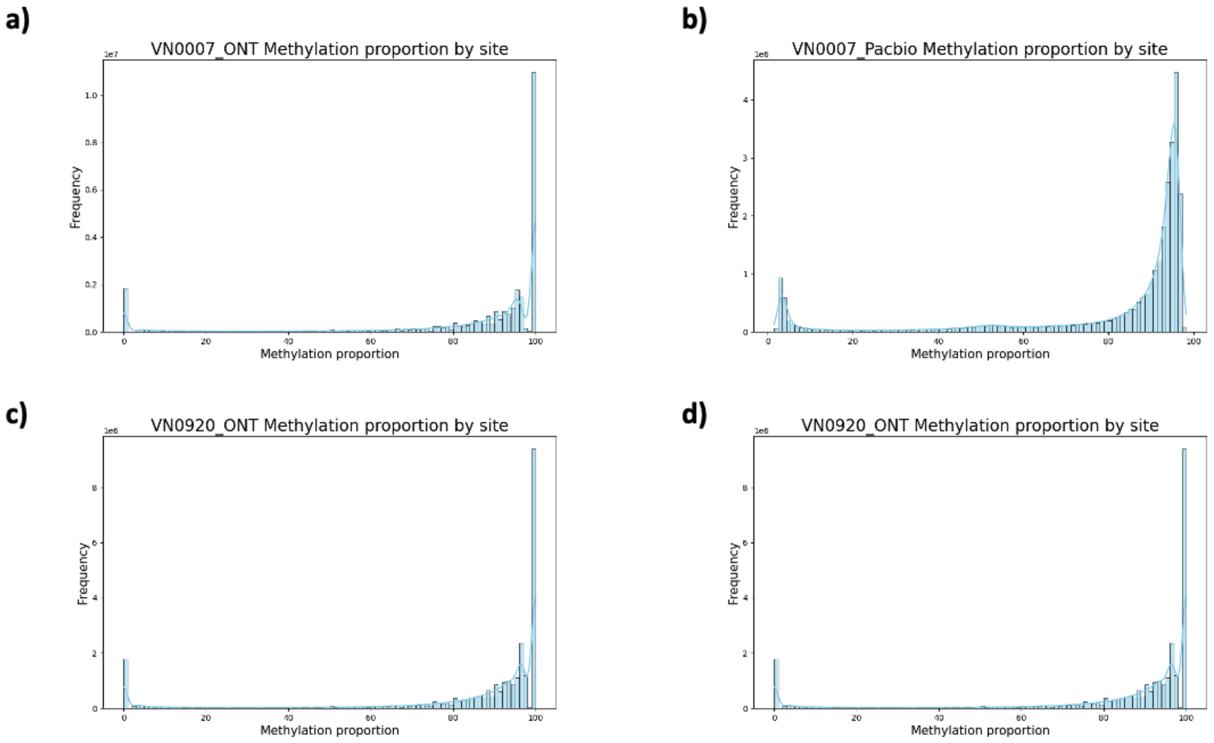


**Supplementary Figure 12.** The plots show distribution of methylation proportions for two Vietnamese samples, comparing results obtained from ONT and Pacbio long-read sequencing.

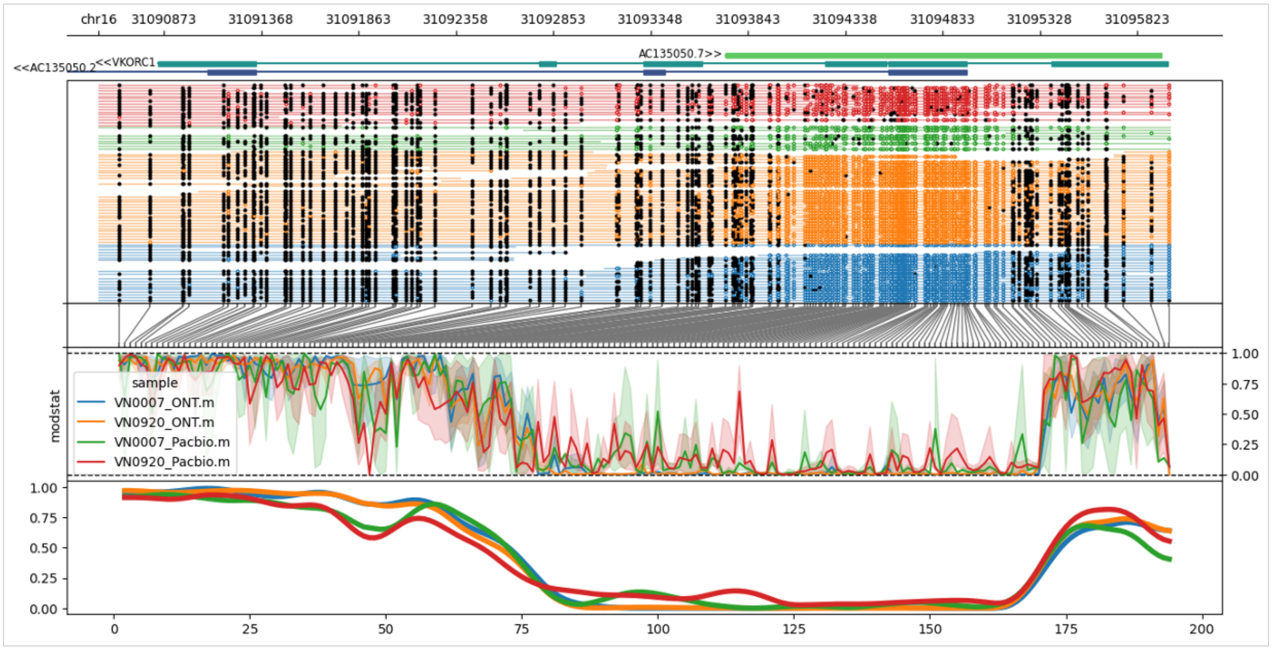


**Supplementary Figure 13.** The figure compares methylation patterns across the VKORC1 gene between two Vietnamese samples with ONT and Pacbio.

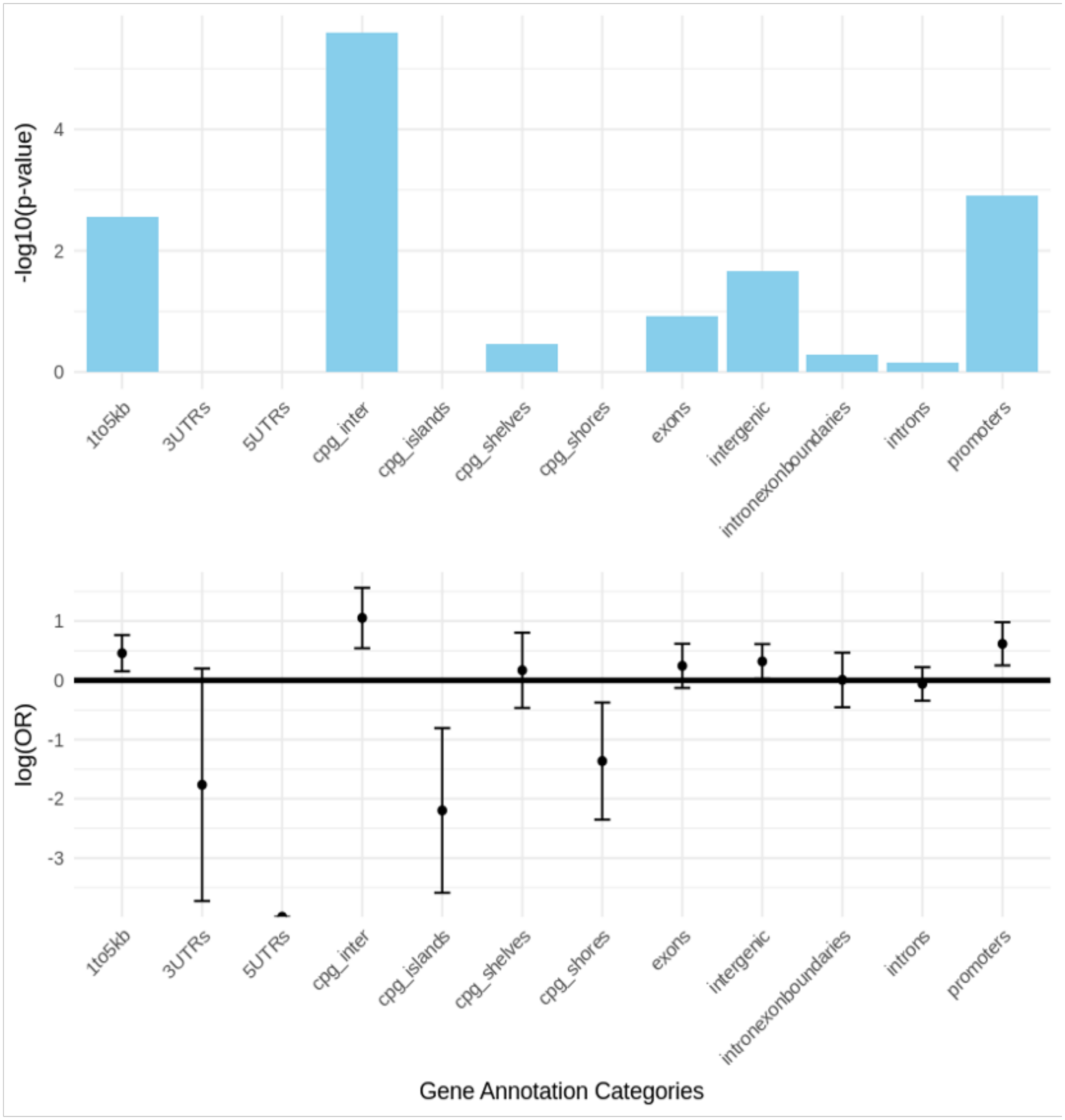


**Supplementary Figure 14.** The enrichment analysis of 202 DMR sites using Fisher's exact test from LOLA package and gene annotation from *annotatr*.
