## Supplementary Tables for "VN1K: a genome graph-based and function-driven multi-omics and phenomics resource for the Vietnamese population"

*This file includes Supplementary Tables 1-4. Additional Supplementary Tables can be found in the Excel file: Supplementary Tables.xlsx*

**Supplementary Table 1**. Statistics of CNV from VN1K. The range of CNVs per individual is also similar, with slight variations. The South shows a slightly higher lower bound (133) compared to the North (128) and Central (130). The ratio between deletions and duplications (DEL/DUP ratio) is consistent across all regions, hovering around 6.9.

| Statistics | Region | | |
|---|---|---|---|
| | **North** | **Central** | **South** |
| Number of samples | 401 | 432 | 178 |
| Average number of CNVs/individual | 149.99 | 150 | 151.55 |
| Median number of CNVs/individual | 150 | 151 | 152 |
| Range of number of CNVs/individual | 128-170 | 130-173 | 133-172 |
| Ratio Between DEL, DUP CNV | 6.96 | 6.92 | 6.95 |

**Supplementary Table 2**. Quality assessment of assembly samples by QUAST. For Vn920, WENGAN-M has fewer contigs but larger contigs and a higher genome fraction compared to WENGAN-A. Both assemblers show similar GC content. WENGAN-M for VN007 again has fewer contigs but larger contigs and a higher genome fraction compared to WENGAN-A. The number of genomic features is significantly higher in WENGAN-M.

| Sample | VN920 | | | | | | | | | | | |
|---|---|---|---|---|---|---|---|---|---|---|---|---|
| Assembler | SR | LR | # contigs (>= 50000 bp) | # contigs | Largest contig | Total length | # misassembled contigs | Genome fraction (%) | # genomic features | N50 | L50 | GC (%) |

| | | | | | | | | | | | | |
|---|---|---|---|---|---|---|---|---|---|---|---|---|
| WENGAN-M | 30.6X | 25X | 995 | 1818 | 51,112,100 | 2,764,468,443 | 386 | 90.706 | 2804950 + 16593 part | 9,674,427 | 82 | 40.86 |
| WENGAN-A | 30.6X | 25X | 940 | 1989 | 37,251,183 | 2,610,241,688 | 366 | 85.671 | 2711537 + 15438 part | 9,252,486 | 88 | 40.94 |
| | VN920 | vnref5 | | | | 3,217,374,765 | | | | | | 41.01 |
| WENGAN-M | 30.6X | 25X | 995 | 1818 | 51,112,100 | 2,764,468,443 | 388 | 90.707 | | 9,674,427 | 82 | 40.86 |
| WENGAN-A | 30.6X | 25X | 940 | 1989 | 37,251,183 | 2,610,241,688 | 367 | 85.673 | | 9,252,486 | 88 | 40.94 |
| Sample | VN007 | | | | | | | | | | | |
| Assembler | SR | LR | # contigs (>= 50000 bp) | # contigs | Largest contig | Total length | # misassembled contigs | Genome fraction (%) | # genomic features | N50 | L50 | GC (%) |
| WENGAN M | 33.4X | 24.8X | 1,097 | 1,986 | 28,636,227 | 2,749,603,437 | 403 | 90.22 | 2800585 + 18308 part | 6,351,475.00 | 127.00 | 40.86 |
| WENGAN-A | 33.4X | 24.8X | 3,249 | 4,653 | 8,098,930 | 2,694,425,134 | 454 | 88.47 | 2752283 + 24303 part | 1,565,648.00 | 509.00 | 40.85 |
| | VN007 | vnref5 | | | | 3,217,374,765 | | | | | | 41.01 |
| WENGAN-M | 33.4X | 24.8X | 1,097 | 1,986 | 28,636,227 | 2,749,603,437 | 404 | 90.22 | | 6,351,475.00 | 127.00 | 40.86 |
| WENGAN-A | 33.4X | 24.8X | 3,249 | 4,653 | 8,098,930 | 2,694,425,134 | 453 | 88.47 | | 1,565,648.00 | 509.00 | 40.85 |

**Supplementary Table 3.** Statistics of quantitative traits from VN1K in three main regions.

| Trait (Unit) | North | | Central | | South | | Total |
|---|---|---|---|---|---|---|---|
| | Male | Female | Male | Female | Male | Female | |
| Age (years) (n=1008) | 38.45 (3.65) | 39.94 (3.85) | 39.33 (3.64) | 40.1 (3.53) | 38.34 (3.15) | 38.47 (3.18) | 38.34(6.99) |
| BMI (n=1008) | 23.35 (2.85) | 21.73 (2.04) | 23.36 (2.89) | 22.21 (2.55) | 22.85 (3.9) | 22.08 (3.02) | 22.62(2.86) |
| Chest size (centimeters) (n=1008) | 89.16 (6.39) | 85.89 (5.01) | 88.94 (6.52) | 87.52 (5.49) | 87.87 (8.91) | 84.51 (7.19) | 87.56(6.51) |
| Diastolic blood pressure (mmHg) (n=1008) | 73.14 (9.21) | 65.28 (8.31) | 73.78 (10.02) | 67.64 (9.34) | 68.63 (9.91) | 63.08 (8.84) | 69.20(10.00) |
| Heartbeat (BPM-beats per minute) (n=1008) | 73.67 (9.49) | 75.1 (7.72) | 74.57 (11.03) | 75.0 (8.55) | 75.19 (11.64) | 76.39 (10.26) | 74.69(9.61) |
| Height (centimeters) (n=1008) | 167.38 (5.93) | 156.05 (4.61) | 166.27 (5.99) | 155.3 (5.13) | 167.78 (5.62) | 155.74 (5.72) | 157.43(25.79) |
| Hip size (centimeters) (n=1008) | 92.8 (6.5) | 88.68 (6.22) | 94.1 (8.1) | 90.83 (6.83) | 95.71 (7.86) | 87.69 (8.31) | 89.36(15.84) |
| Systolic blood pressure (mmHg) (n=1008) | 117.48 (11.8) | 105.66 (10.24) | 120.27 (13.07) | 110.86 (11.81) | 116.84 (13.62) | 107.73 (10.8) | 113.29(13.07) |
| Weight (kilograms) (n=1008) | 66.36 (8.76) | 52.9 (5.32) | 65.34 (8.37) | 53.55 (6.47) | 67.11 (9.05) | 53.59 (8.07) | 58.15(13.34) |
| ALT (U/L) (n=977) | 29.97 (21.51) | 17.85 (20.1) | 28.23 (15.93) | 14.23 (5.25) | 28.73 (14.74) | 17.57 (12.58) | 23.32(17.83) |
| AST (U/L) (n=977) | 26.67 (13.03) | 21.61 (14.35) | 26.51 (7.61) | 20.1 (3.71) | 25.84 (6.72) | 21.59 (6.56) | 24.13(10.33) |
| Acid Uric (µmol/L) (n=892) | 410.21 (74.14) | 281.07 (53.09) | 392.23 (69.43) | 271.95 (56.6) | 377.38 (74.31) | 281.58 (52.35) | 339.78(88.03) |
| WBC (G/L) (n=1006) | 6.68 (1.83) | 6.14 (1.55) | 6.54 (1.59) | 6.16 (1.68) | 6.77 (1.59) | 6.32 (1.72) | 6.49(1.70) |
| Creatinine (µmol/L) (n=977) | 82.98 (10.52) | 61.08 (10.29) | 86.65 (11.46) | 63.69 (9.36) | 88.41 (11.76) | 66.75 (8.57) | 74.49(15.13) |
| Glucose (mmol/L)) (n=977) | 5.18 (0.77) | 4.92 (0.51) | 5.3 (0.89) | 4.97 (0.64) | 5.14 (0.39) | 5.11 (0.72) | 5.08(0.67) |
| HDL-C (mmol/L) (n=994) | 1.19 (0.27) | 1.4 (0.28) | 1.18 (0.25) | 1.41 (0.27) | 1.22 (0.25) | 1.36 (0.25) | 1.29(0.29) |
| HbA1c (%) (n=915) | 5.57 (0.57) | 5.4 (0.37) | 5.47 (0.53) | 5.42 (0.48) | 5.26 (0.39) | 5.38 (0.49) | 5.44(0.49) |

| | | | | | | | |
|---|---|---|---|---|---|---|---|
| LDL-C (mmol/L) (n=894) | 3.42 (0.69) | 2.99 (0.6) | 3.46 (0.68) | 3.08 (0.62) | 3.42 (0.65) | 3.19 (0.63) | 3.24(0.68) |
| Total Cholesterol (mmol/L) (n=898) | 5.07 (0.86) | 4.72 (0.77) | 5.35 (0.98) | 4.95 (0.8) | 5.23 (0.77) | 5.02 (0.73) | 5.03(0.88) |
| Triglyceride (mmol/L) (n=896) | 1.87 (1.17) | 1.12 (0.84) | 2.27 (2.21) | 1.13 (0.67) | 1.59 (0.78) | 1.12 (0.67) | 1.58(1.46) |
| Ure (mmol/L) (n=965) | 5.22 (1.05) | 4.39 (0.99) | 5.09 (1.15) | 4.29 (1.02) | 4.83 (1.16) | 4.29 (0.99) | 4.71(1.12) |

**Supplementary Table 4.** The frequencies of KIR genes calling by KPI for VN1K and 1kGP-HC

| **Gene** | **1kGP-HC** | **VN1K** | **Fisher's exact test p-value** |
|---|---|---|---|
| 2DL1 | 0.98 | 0.98 | 1.00 |
| 2DL2 | 0.49 | 0.33 | 0.00 |
| 2DL3 | 0.90 | 0.97 | 0.00 |
| 2DL4 | 1.00 | 1.00 | NA |
| 2DL5 | 0.53 | 0.45 | 0.00 |
| 2DP1 | 0.98 | 0.98 | 1.00 |
| 2DS1 | 0.39 | 0.38 | 0.59 |
| 2DS2 | 0.47 | 0.33 | 0.00 |
| 2DS3 | 0.28 | 0.23 | 0.00 |
| 2DS4 | 0.94 | 0.96 | 0.02 |
| 2DS5 | 0.36 | 0.27 | 0.00 |
| 3DL1 | 0.94 | 0.96 | 0.02 |
| 3DL2 | 1.00 | 1.00 | NA |
| 3DL3 | 1.00 | 1.00 | NA |
| 3DP1 | 1.00 | 1.00 | NA |
| 3DS1 | 0.35 | 0.39 | 0.03 |
